# A complex osteoporotic milieu is associated with arterial stiffening and PDGF-BB-mediated calcification of human smooth muscle cells

**DOI:** 10.1101/2025.10.25.684499

**Authors:** Wera Pustlauk, Toralf Roch, Ioanna Maria Dimitriou, Arturo Blázquez Navarro, Ulrik Stervbo, Moritz Anft, Julian Braun, Rainer Wirth, Maryam Pourhassan, Bjoern Buehring, Doruk Akgün, Patrizia Wehler, Sarah Skrzypczyk, Gerald Grütz, Sven Rahmann, Felix Sebastian Seibert, Timm Henning Westhoff, Nina Babel, Sven Geißler

**Affiliations:** Berlin Institute of Health Center for Regenerative Therapies (BCRT), Charité - Universitätsmedizin Berlin, corporate member of Freie Universität Berlin, Humboldt-Universität zu Berlin, and Berlin Institute of Health, Berlin, Germany; Julius Wolff Institute for Biomechanics and Musculoskeletal Regeneration, Berlin Institute of Health at Charité - Universitätsmedizin Berlin, Berlin, Germany; Center for Translational Medicine, Department of Internal Medicine I, Marien Hospital Herne - University Hospital of the Ruhr-Universität Bochum, Germany; DZHK (German Centre for Cardiovascular Research), partner site Berlin, Germany; CheckImmune GmbH, Berlin, Germany; Department of Geriatric Medicine, Marien Hospital Herne, Ruhr-Universität Bochum, Germany; Rheumazentrum Ruhrgebiet, Ruhr-Universität Bochum, Germany; Bergisches Rheuma-Zentrum Wuppertal, Germany; Center for Musculoskeletal Surgery, Department for Shoulder and Elbow Surgery, Charité - Universitätsmedizin Berlin, Germany; Algorithmic Bioinformatics, Center for Bioinformatics Saar, Saarland Informatics Campus, Saarland University, Saarbrücken, Germany; Medical Department 1, Marien Hospital Herne - University Hospital of the Ruhr-Universität Bochum, Germany

## Abstract

Accumulating evidence links skeletal and vascular aging, yet the pathological cross-talk between osteoporotic bone and vascular calcification remains insufficiently understood. In this retrospective exploratory study, we show that osteoporotic individuals exhibit a distinct systemic milieu characterized by elevated eosinophil, neutrophil, and platelet counts, as well as altered levels of sCD40L, PDGF-BB, osteopontin, and SDF-1. These changes correlate with both bone metabolism and arterial stiffness, indicating a multifactorial bone-vascular interplay. Moreover, osteoporotic sera accelerated calcification of human vascular smooth muscle cells *in vitro*, with PDGF-BB emerging as a central mediator. Inhibition of PDGF-BB downstream signaling blocked this effect, suggesting a mechanistic role of PDGF-BB in linking the osteoporotic milieu to vascular calcification. Conceptualizing this cross-talk as a complex adaptive system advances our understanding of the disease dynamics between localized bone loss and vascular pathologies. Furthermore, it may guide therapeutic strategies, with PDGF-BB as a potential target, to support healthier aging.

## Introduction

Increasing clinical and epidemiological evidence demonstrates a distinct association between osteoporosis and vascular calcification.^1–4^ Independent of shared risk factors such as advanced age, sedentary lifestyle, smoking, menopause, and diabetes,^5–7^ the pathological pathways linking bone and vasculature remain under investigation.^8^ Recent murine studies have identified platelet-derived growth factor beta (PDGF-BB), secreted by preosteoclasts in the bone marrow, as a key mediator of arterial stiffening and vascular calcification in the brain.^9–11^ Earlier studies, moreover, established matrix Gla protein as an important paracrine inhibitor of calcification in both bone and vasculature in mice and human atherosclerotic lesions.^7,12^ Furthermore, osteopontin (OPN) is secreted by activated osteoclasts and promotes mineralization in human bone,^13^ yet it has also been reported to inhibit arterial calcification.^7,14^

As osteoporosis, atherosclerosis, and vascular calcification are driven by inflammatory processes, an inflammatory link between both conditions has also been suggested.^15,16^ Atherosclerotic plaque development involves extensive infiltration of the arterial wall by macrophages, T cells, and B cells, accompanied by the release of cytokines including monocyte chemoattractant protein-1 (MCP-1), interferon-γ (IFN-γ), and interleukin (IL)-1.^17^ By contrast, osteoporosis is associated with increased osteoclast formation from macrophages,^18,19^ with the osteoprotegerin (OPG)/receptor activator of NF-кB (RANK)/RANK ligand (RANKL) system serving as a key regulator of this process in bone.^20^ Murine studies investigating estrogen depletion revealed that RANKL-OPG signaling suppresses the expression of pro-inflammatory cytokines, such as macrophage colony-stimulating factor, IL-1, IL-6, and tumor necrosis factor-α (TNF-α),^21,22^ suggesting that these factors may act as systemic drivers of vascular calcification in osteoporotic women. However, TNF-α and IL-6 have thus far been reported as mediators of vascular calcification only in hemodialysis patients, while data from other cohorts are lacking.^23^

The numerous factors investigated in previous studies point to a multifaceted interplay between bone and vasculature. The theory of complex adaptive systems (CAS) conceptualizes such interplays as dynamic, interdependent interactions among diverse components capable of self-organization.^24–27^ In response to their environment, such systems and their subsystems can learn from experience, adapt rules, and thus new behavioral patterns and structures emerge.^25,26^ Thus, small changes in one component can cause distinct systemic effects.^26,27^ Applying CAS theory to the pathological bone-vascular cross-talk could therefore advance our understanding of disease dynamics and help to identify novel intervention targets.^24,27^

In this hypothesis-generating human study, we retrospectively explore the inflammatory status of individuals with different levels of bone mineral density (BMD). By combining clinical data analysis with *ex vivo* screening of blood and serum samples, we identify a systemic osteoporotic milieu associated with arterial stiffening. In a mechanistic *in vitro* study we then investigated the role of PDGF-BB in vascular calcification in osteoporotic patients, extending murine findings to humans. Finally, we conceptualize the pathological bone-vascular cross-talk as a CAS to improve understanding of disease complexity and help to identify novel opportunities for targeted interventions that support healthier aging.

## Results

### Study groups and their characteristics indicate links between osteoporosis, frailty, and arterial stiffening

To explore the interplay between osteoporotic bone and vasculature, we retrospectively analyzed 100 participants (23 - 91 years; 54 women, 46 men) recruited at Ruhr-Universität Bochum and Charité - Universitätsmedizin Berlin. Wherever possible, we ensured comparable sex distribution across geriatric study groups from the Ruhr-Universität Bochum and non-geriatric study groups recruited for Charité - Universitätsmedizin Berlin as well as availability of diagnostic data and biosamples. Participants were first classified by bone mineral density (BMD) into osteoporotic (OPO), osteopenic (OPE), or non-osteoporotic controls (CON), based on T-scores or, if not available, by documented self-report based on prior medical examinations (Fig. 1 A, Table 1, Extended Data Tables 1 and 2). To reflect the heterogeneity of the older population and address potential biases of age and frailty within the non-osteoporotic controls, we distinguished three subgroups: middle-aged healthy controls (CON-M, n=11, 40.8 ± 12.4 years), older healthy controls (CON-H, n=19, 72.3 ± 5.5 years), and older frail controls (CON-F, n=16, 65.2 ± 10.3 years).

**Fig. 1:**
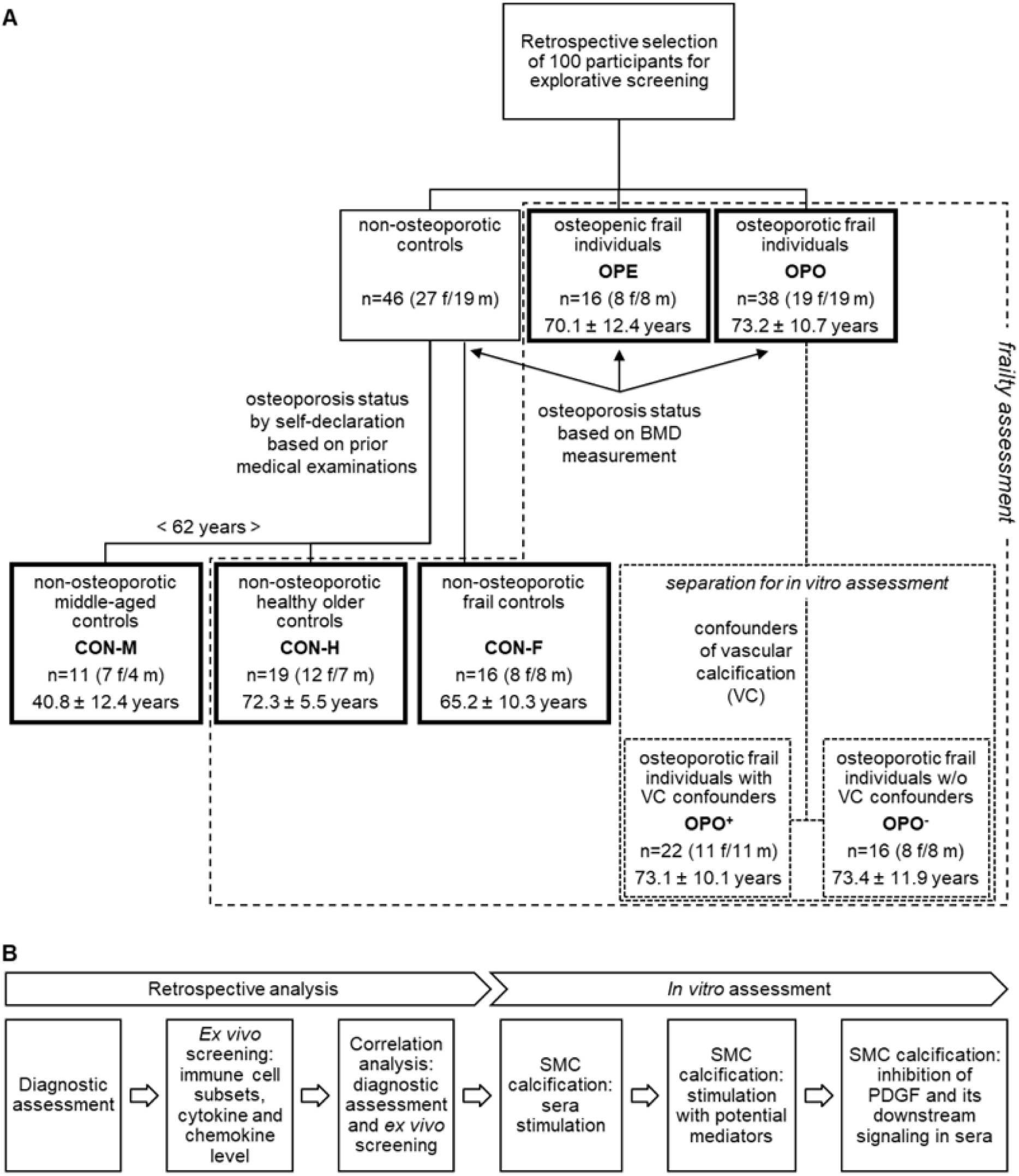
Study groups and study outline. **(A)** Study participants were retrospectively stratified according to osteoporosis status, age, and age-related frailty (dotted line; pre-existing orthopedic conditions, osteochondrosis and disc prolapses, rheumatoid arthritis, number of falls within the last year, difficulty climbing stairs). For *in vitro* assessments, subgroups of osteoporotic individuals with or without confounders of vascular calcification (diabetes, use of blood sugar reducing medication, hyperuricemia, or renal insufficiency) were analyszed separately. BMD - bone mineral density, f - female, m - male, SMC - smooth muscle cells, VC - vascular calcification. **(B)** Consecutive study outline.

**Table 1:**
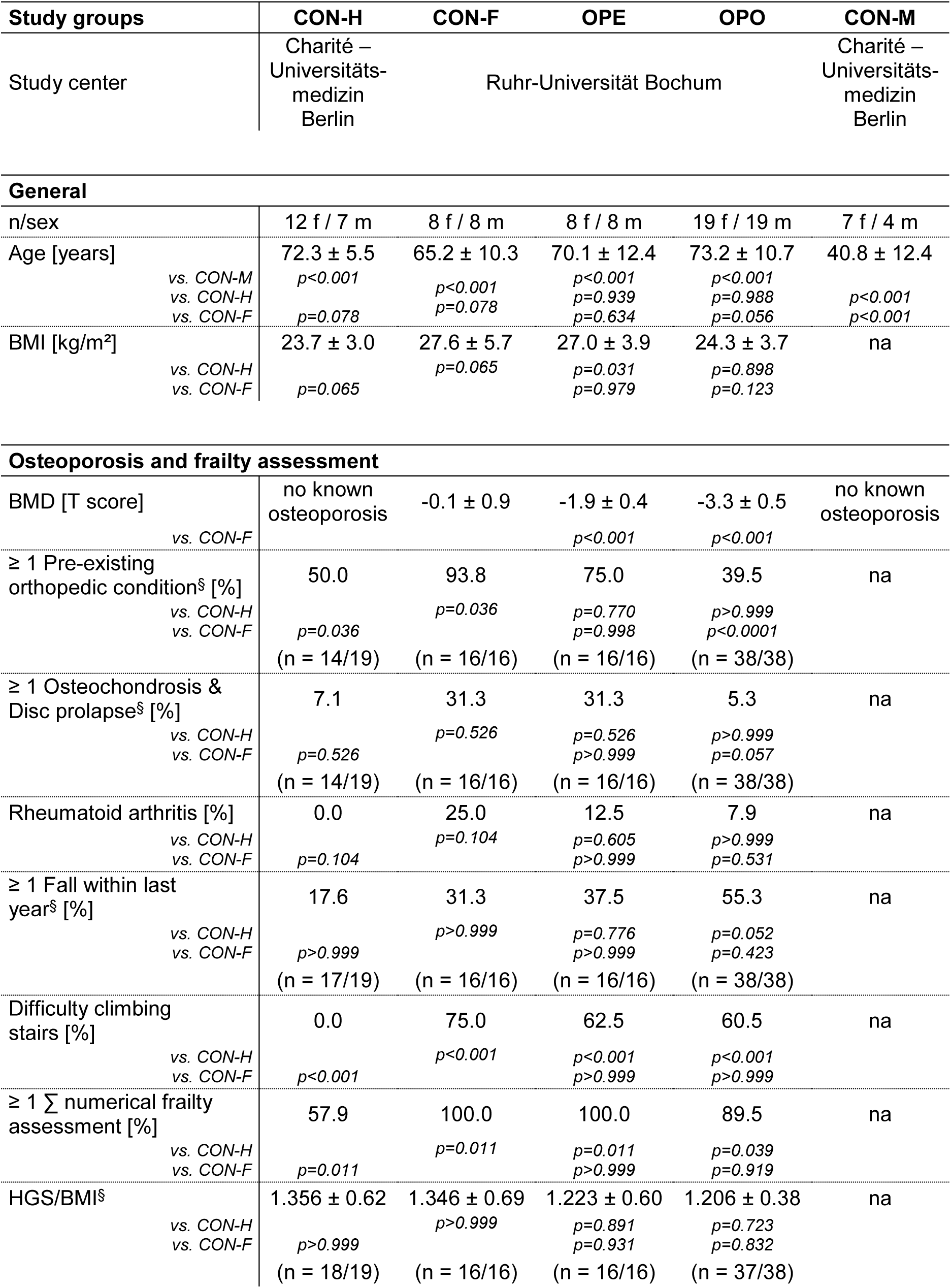
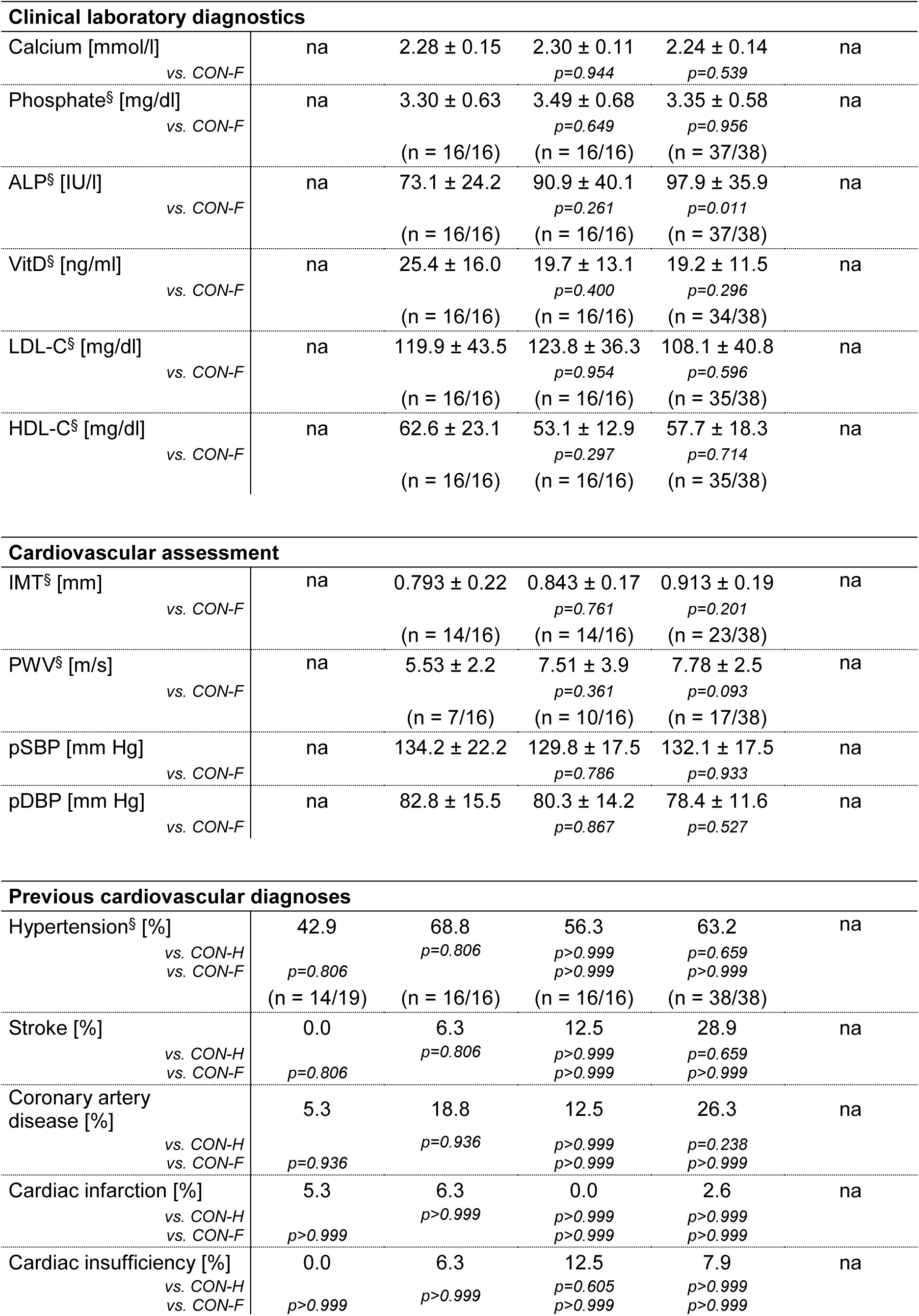
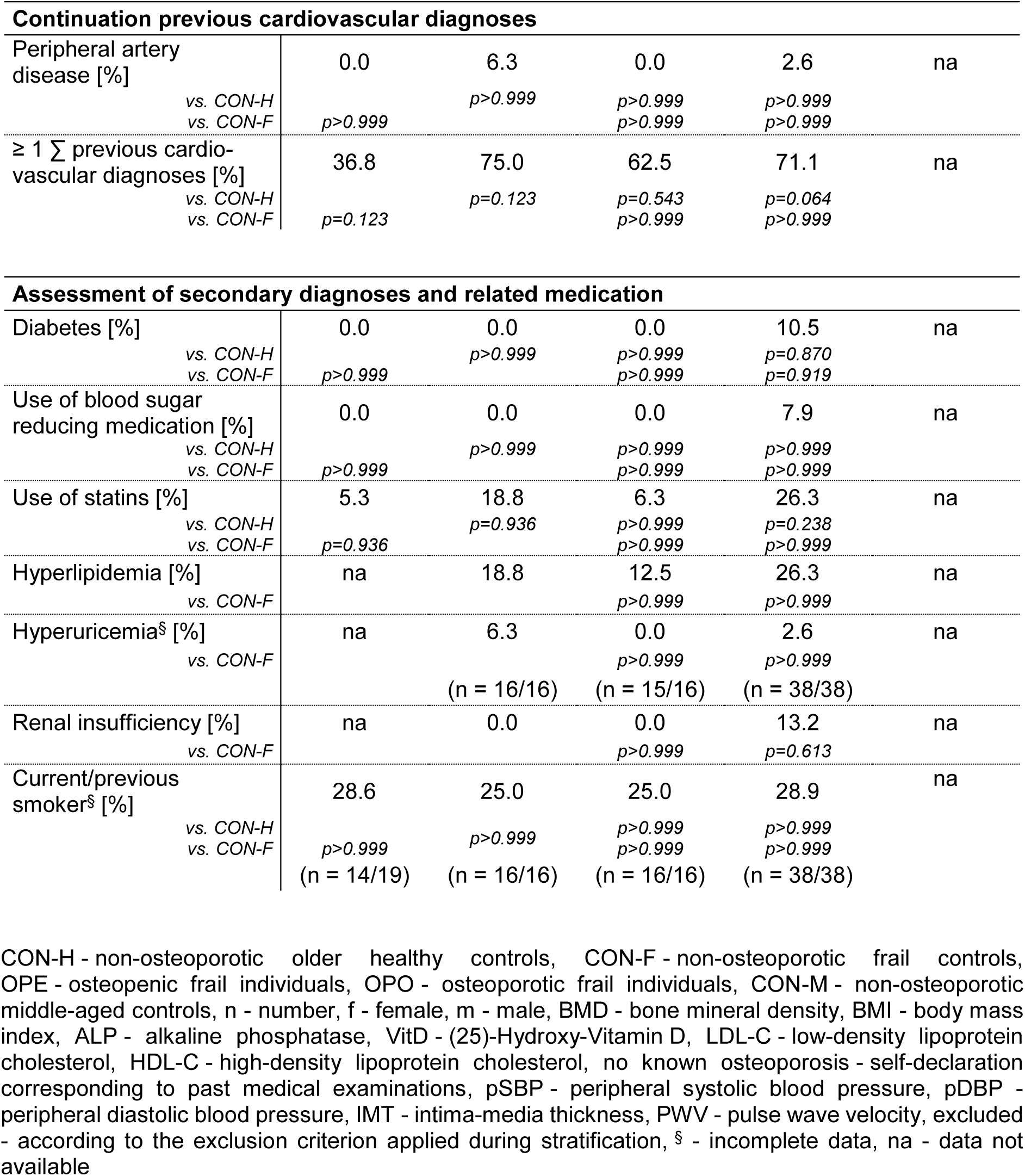
Diagnostic data of non-osteoporotic, osteopenic, and osteoporotic individuals. Data are given as n, mean ± standard deviation, overall percentages, or percentages of individuals presenting with more than one condition. Differences between study groups were analyzed using a Brown-Forsythe and Welch ANOVA with Dunnett’s post-test for numerical data or Fisher’s exact test with Bonferroni correction for binary variables (p<0.05 vs. CON-M, CON-H, or CON-F).

Frailty assessments were conducted in all older participants. However, since the recorded items differed between study centers,^28–30^ comparative analysis included only data collected in a comparable manner (number of falls within the last year, difficulty climbing stairs, and hand grip strength (HGS)), complemented by available comorbidity data (pre-existing orthopedic conditions, osteochondrosis and disc prolapses, rheumatoid arthritis). This analysis revealed that geriatric OPE (n=16, 70.1 ± 12.4 years) and OPO (n=38, 73.2 ± 10.7 years) participants had significantly higher frailty scores and markedly greater difficulty climbing stairs than CON-H, recruited at retirement homes. However, their frailty status was comparable to CON-F. Additionally, OPO individuals experienced considerably more falls in the past year. All groups provided data and biosamples for retrospective *ex vivo* analysis, followed by mechanistic *in vitro* studies (overview: Fig. 1 B). To control for confounders of vascular calcification in the *in vitro* experiments, OPO individuals were further stratified into OPO^+^ (with diabetes, hyperuricemia, or renal insufficiency; n=22, 73.1 ± 10.1 years) and OPO^-^ (without these comorbidities; n=16, 73.4 ± 11.9 years).

Additional analysis of diagnostic data revealed significantly higher systemic alkaline phosphatase (ALP) levels, a marker of bone turnover, in osteoporotic individuals (OPO) compared with frail controls (CON-F). Simultaneously, osteoporotic individuals (OPO) showed trends of increased intima-media thickness (IMT) and higher pulse wave velocity (PWV), both indicators of arterial stiffness. No differences were detected in lipid profiles (LDL-C, HDL-C) or peripheral systolic and diastolic blood pressure (pSBP, pDBP) among study groups. Results remained consistent after re-analysis stratified by sex (Extended Table 1 and 2), except for the numerical frailty assessment, which was significantly higher in men but not in woman, likely due to the higher prevalence of pre-existing orthopedic conditions in healthy female controls.

### Low bone mineral density is associated with a systemic multifactorial osteoporotic milieu

To investigate cellular changes associated with low BMD while controlling for the confounding effects of age and frailty, we analyzed peripheral immune cell subsets available from flow cytometry analysis in CON-F, OPE, and OPO (gating strategy in Supplementary Fig. 1). Monocyte, T cell, natural killer (NK) cell, or B cell numbers did not differ between the groups (Extended Data Fig. 1). In contrast, granulocyte, eosinophil, neutrophil, and platelet counts were significantly higher in OPO (Fig. 2 A-E). Correspondingly, inflammatory ratios such as granulocyte-to-lymphocyte, basophil-to-lymphocyte, eosinophil-to-lymphocyte (ELR), and neutrophil-to-lymphocyte ratios (NLR), as well as the systemic inflammation index (SII; NLR × platelet count),^31^ were elevated in OPO individuals compared with CON-F (Extended Data Table 3). Sex-stratified analysis showed that eosinophils, neutrophils, ELR, and SII were predominantly elevated in male OPO individuals (Extended Data Fig. 2, Extended Data Table 4), whereas platelet counts and NLRs were elevated in both sexes, indicating a generally altered systemic immune cell status.

**Fig. 2:**
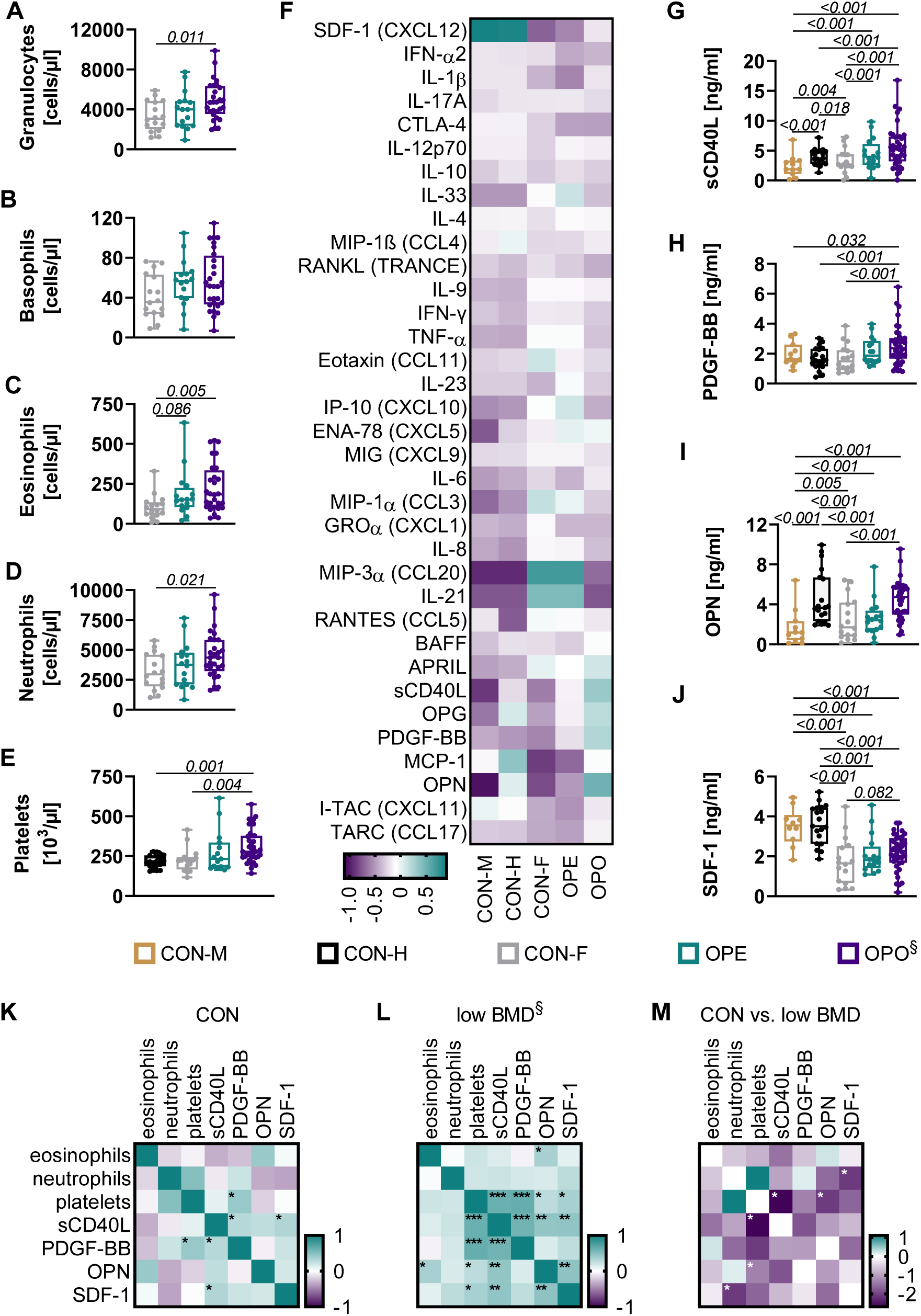
Osteoporosis is associated with higher eosinophil, neutrophil, and platelet counts, as well as higher sCD40L and PDGF-BB levels. Flow cytometry analysis of **(A)** total granulocytes, **(B)** basophils, **(C)** eosinophils, **(D)** neutrophils, and platelets in the peripheral blood of non-osteoporotic frail controls (CON-F; n=16), osteopenic frail individuals (OPE; n=16), and osteoporotic frail individuals (OPO; n=28 out of 38). Differences in immune cell subsets were assessed using a Brown-Forsythe and Welch ANOVA with Dunnett’s T3 multiple comparisons test vs. CON-F if data were normally distributed. Non-normal data were analyzed with a Kruskal-Wallis test and Dunn’s multiple comparison. ^§^ indicates incomplete immune cell subset data. **(E)** Sera screening of non-osteoporotic middle-aged controls (CON-M), non-osteoporotic healthy older controls (CON-H), CON-F, OPE, and OPO individuals using bead-based multiplex assays. Clustered expression levels are given as median Z-scores. Factors with significantly different expression levels or distinct trends with p<0.1 in OPE or OPO individuals vs. CON-M, CON-H, and CON-F include **(G)** sCD40L, **(H)** PDGF-BB, **(I)** OPN, and **(J)** SDF-1. Differences between serum factors were assessed using two-way ANOVAs with Dunnett’s post-test (p<0.05) vs. CON-H, CON-M, or CON-F. Only significant differences (p<0.05) and distinct trends with p<0.1 are shown. **(K)** Spearman correlation of the significantly different cellular and serum factors in non-osteoporotic controls (CON: CON-M, CON-H, CON-F) and **(L)** in osteopenic and osteoporotic individuals (low BMD: OPE, OPO). **(M)** Differences between correlation coefficients of CON and low BMD were assessed using Fisher’s Z-test; * p<0.05, ** p<0.001, *** p<0.0001.

Multiplex assays further revealed distinct cytokine and chemokine profiles in frail individuals (CON-F, OPE, OPO) compared to middle-aged (CON-M) and healthy older (CON-H) controls (Fig. 2 F). Specifically, CON-F, OPE, and OPO individuals showed reduced SDF-1 levels and minor increases of IL-6, IL-8, IL-21, MIP-1α, MIP-3α, and IP-10 (CXCL10), consistent with a low-grade pro-inflammatory milieu.

Interestingly, classical pro-inflammatory cytokines and chemokines, including TNFα, IFN-γ, IL-1β, IL-6, and IL-17A, were not significantly altered in OPE and OPO individuals compared to controls (CON-M, CON-H, CON-F; total and sex-separated analysis: p>0.9999; Fig. 2 F, Supplementary Fig. 4 A). However, sCD40L and PDGF-BB levels were significantly higher in OPO sera compared with controls and across sexes (Fig. 2 G & H, Extended Data Fig. 3 B). By contrast, OPN (Fig. 2 I) and SDF-1 (Fig. 2 J) levels differed mainly among control subgroups, with both factors being significantly lower in frail than in healthy individuals.

Correlation analysis highlighted a particularly strong interrelation between sCD40L, PDGF-BB, and platelet counts (Spearman ρ > 0.50) in individuals with low BMD (Fig. 2 K-M), suggesting an interconnection between these factors with progressive osteoporosis. Similar strong correlations between these three factors were confirmed in an exploratory sex-segregated analysis (Extended Data Fig. 3 C & D), but were more pronounced in male OPO individuals. Additional significant associations were observed in individuals with low BMD (Fig. 2L), namely between OPN and sCD40L, eosinophils, or platelets, as well as between SDF-1 and sCD40L, OPN, or platelets.

These distinct correlations were either absent or less pronounced in controls (Fig. 2 K-M), supporting the hypothesis of a multifactorial interdependent interplay in osteoporotic patients. Together, these findings indicate that low BMD is characterized by a systemic milieu of elevated granulocyte and platelet counts, higher sCD40L and PDGF-BB levels, and altered OPN and SDF-1 levels. This signature differs from the frailty-associated low-grade inflammation and appears to be particularly pronounced in osteoporotic men.

### The systemic osteoporotic milieu is associated with arterial stiffening

To assess potential associations between systemic alterations, osteoporosis, and the cardiovascular risk, we performed correlation analyses of clinical parameters and *ex vivo* screening data. In non-osteoporotic controls, BMD correlated with eosinophil counts and ELR, while ALP showed marked associations with sCD40L and OPN (Table 2). In individuals with low BMD (OPE, OPO), these associations were attenuated or lost and new correlations emerged, such as between SII and BMD and between ELR and ALP. Notably, the correlation coefficients for ALP-eosinophils and ALP-OPN differed significantly between non-osteoporotic controls and individuals with low BMD, suggesting a shift in immune-bone marker relationships in osteoporosis.

**Table 2:**
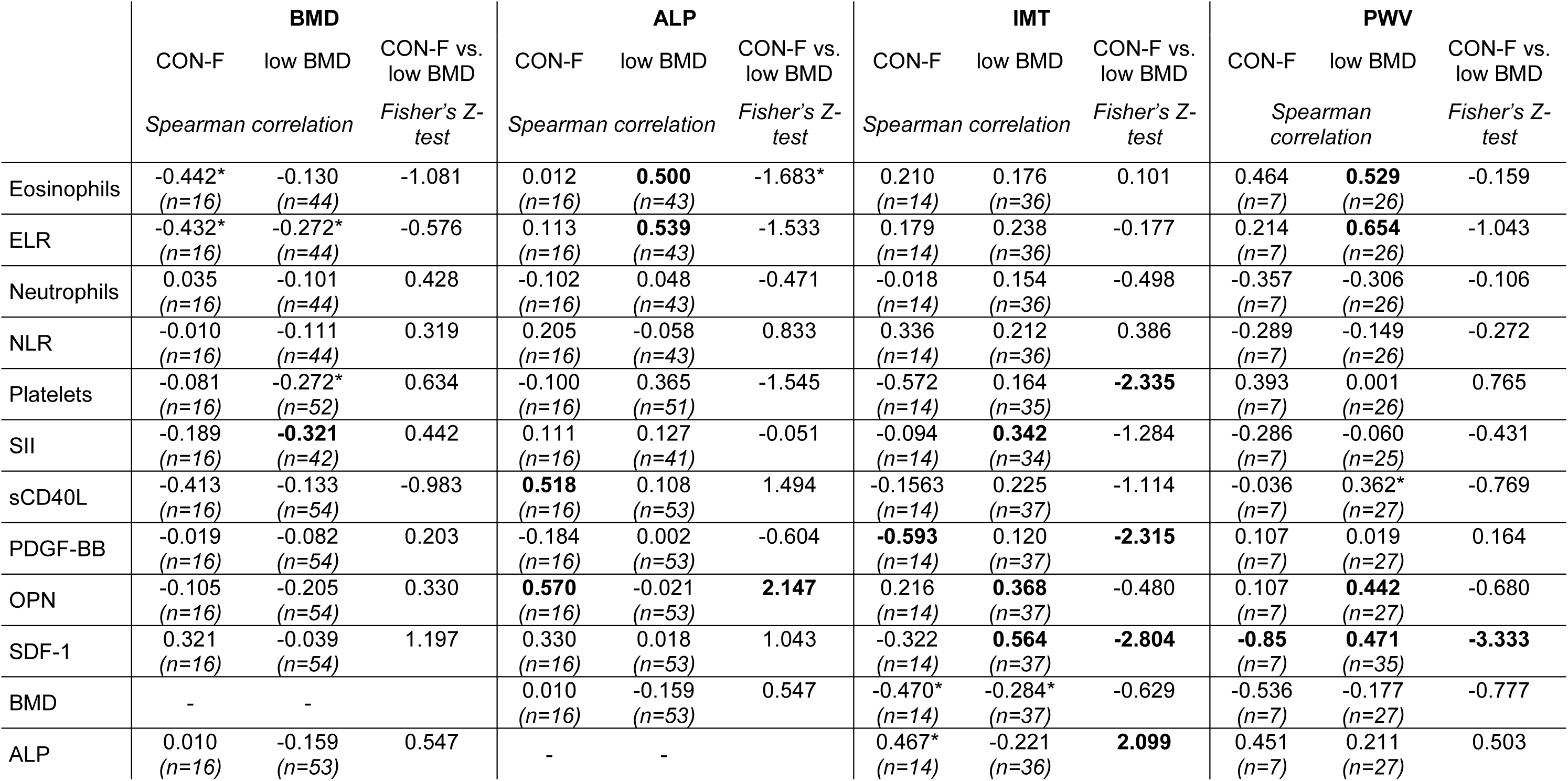
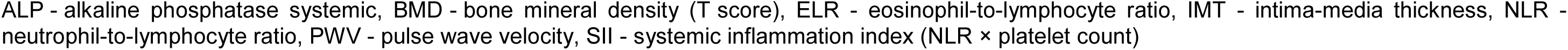
Systemic changes in osteoporosis are associated with arterial stiffening. Spearman correlation of differential immune cell, cytokine, and chemokine levels with diagnostic data of bone turnover and arterial stiffening in non-osteoporotic frail controls (CON-F; total n=16), osteopenic frail individuals (OPE), and osteoporotic frail individuals (OPO) from the Ruhr-Universität Bochum study center. Individuals with low BMD (OPE, OPO; total n=54) were combined for the analysis. Differences between correlation coefficients of CON-F and low BMD were assessed using Fisher’s Z-test. Significant correlations and differences between correlation coefficients with p<0.05 are highlighted in bold; * indicates correlations and differences between correlation coefficients showing distinct trends (p<0.1).

Factors of the osteoporotic milieu also showed altered correlations with markers of vascular stiffness. In individuals with low BMD, IMT correlated with SII, OPN, and SDF-1, while PWV correlated with eosinophils, ELR, sCD40L, and OPN. These patterns were absent in non-osteoporotic controls. Furthermore, correlations between platelets, PDGF-BB, SDF-1, ALP, and IMT, as well as between SDF-1 and PWV differed significantly by BMD. Together, these results indicate that the osteoporotic milieu is characterized not only by disrupted correlations with markers of bone turnover, but also by newly emerging links to vascular stiffness. Therefore, immune cell changes, platelet activation, and altered cytokine/growth factor levels seem to be interconnected with both osteoporosis progression and arterial stiffening.

### Osteoporotic and osteopenic sera induce earlier and stronger smooth muscle cell calcification

To test whether patient sera impact vascular calcification, we exposed primary human smooth muscle cells (SMC) to sera from non-osteoporotic, osteopenic, and osteoporotic individuals using an established human *in vitro* assay that incorporates clinically relevant donor variability.^32^ Sex-mixed sera pools from OPE and OPO individuals induced earlier calcification of SMC and significantly increased deposition of calcified matrix compared to healthy, non-frail individuals of the same age (CON-H; Fig. 3 A-D). No differences were observed between osteoporotic sera from participants with (OPO^+^) or without (OPO^-^) confounders of vascular calcification. This effect was consistent across sexes, with the same trend observed between female and male osteoporotic sera pools (OPO^+^) and their respective controls (Fig. 3 C).

**Fig. 3:**
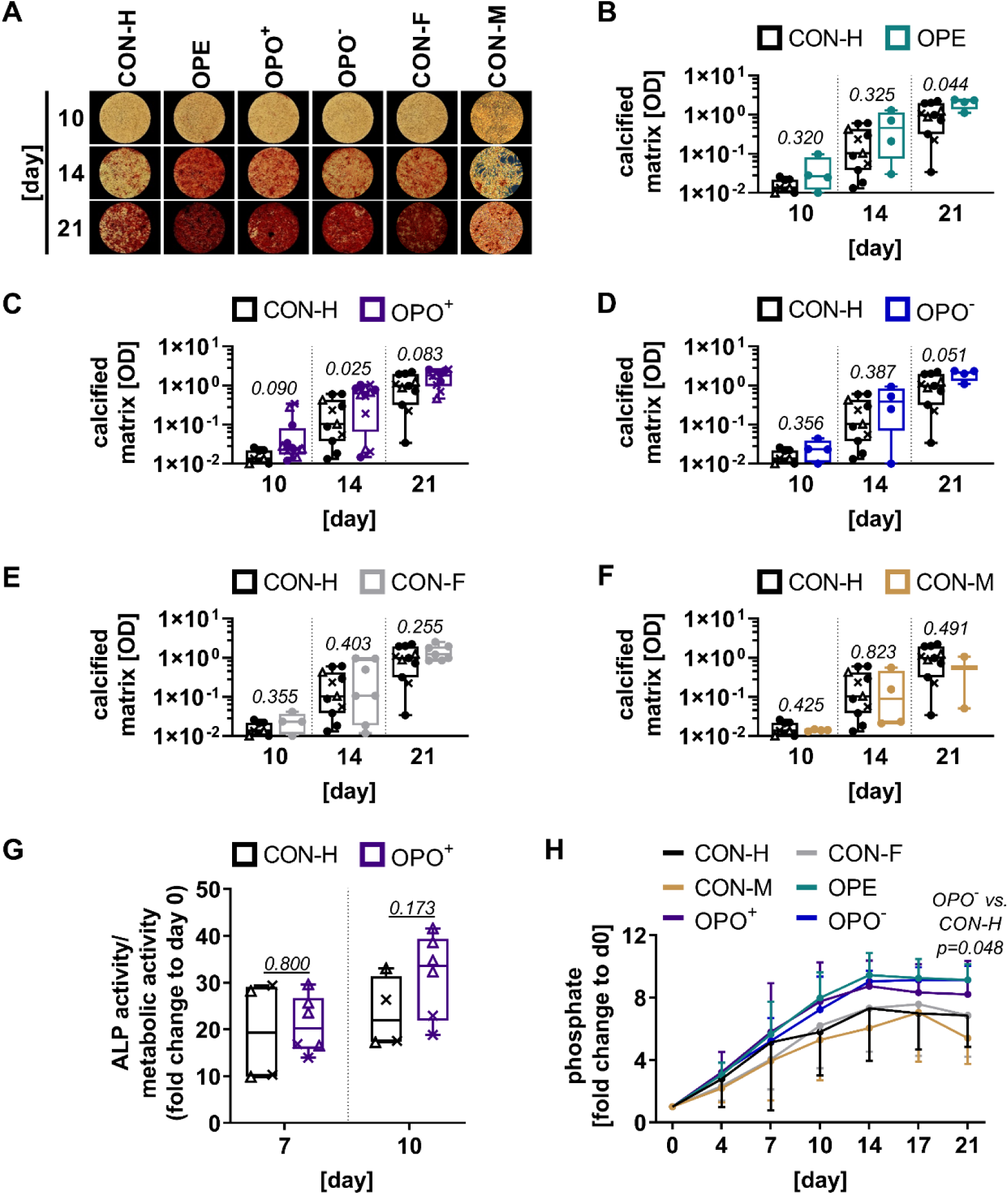
Osteoporotic and osteopenic sera increase SMC calcification. SMC undergoing osteogenic differentiation in an *in vitro* model for SMC calcification were stimulated with sera pools from non-osteoporotic healthy older controls (CON-H), osteopenic frail individuals (OPE), osteoporotic frail individuals with (OPO^+^) or without confounders of vascular calcification (OPO^-^), non-osteoporotic frail controls (CON-F), and non-osteoporotic middle-aged controls (CON-M). Deposited calcified matrix was stained with alizarin red. **(A**) Representative images of deposited calcified matrix and **(B-F)** quantified staining. Differences in SMC calcification at each time point were assessed using an unpaired, two-tailed t-test with Welch’s correction (p<0.05). **(G)** Alkaline phosphatase (ALP) activity of SMC and **(H)** phosphate levels in the supernatant were measured after osteogenic stimulation with sera pools. ALP activity was normalized to the metabolic activity. Differences in ALP activity were assessed using an unpaired, two-tailed t-test with Welch’s correction (p<0.05) for each time point. Phosphate levels at each time point were analyzed using a mixed-effects model with Geisser-Greenhouse correction and Dunnett’s post-test vs. CON-H (p<0.05); only significant differences are shown. Symbols indicate sex-specification of serum pools: ● sex-mixed serum pool, **Δ** female serum pool, **×** male serum pool.

Interestingly, CON-F sera also induced modestly increased SMC calcification compared to CON-H (Fig. 3 E), whereas no differences were observed between CON-H and CON-M sera (Fig. 3 F). This suggests an additional effect of frailty on SMC mineralization *in vitro*.

Increased calcification was accompanied by elevated cellular ALP activity and higher phosphate levels in cultures treated with OPO^+^ compared to CON-H at day ten (Fig. 3 G and H). Similarly elevated phosphate levels were observed in cultures treated with sera from OPE and OPO^-^ individuals. Overall, our findings indicate that sera from osteoporotic and osteopenic individuals promote SMC calcification.

### PDGF-BB increases SMC calcification

We next investigated whether the identified serum mediators, sCD40L, PDGF-BB, OPN, and SDF-1α, directly contribute to increased SMC calcification. Only PDGF-BB, alone or in combination with other factors, significantly increased SMC calcification compared to osteogenic medium (OM) controls (Fig. 4 A-E, Extended Data Fig. 4 A-H). Consistently, these cultures exhibited elevated phosphate levels in the supernatant (Fig. 4 F). By contrast, sCD40L had no effect on phosphate levels in cultures, whereas OPN and SDF-1α treated cultures showed reduced levels compared to OM control. Respectively, combinations of PDGF-BB with OPN and/or SDF-1α resulted in phosphate levels comparable to those of the OM control (Extended Data Fig. 4 I), suggesting a modulatory effect of OPN and SDF-1α on the PDGF-BB-induced increase.

**Fig. 4:**
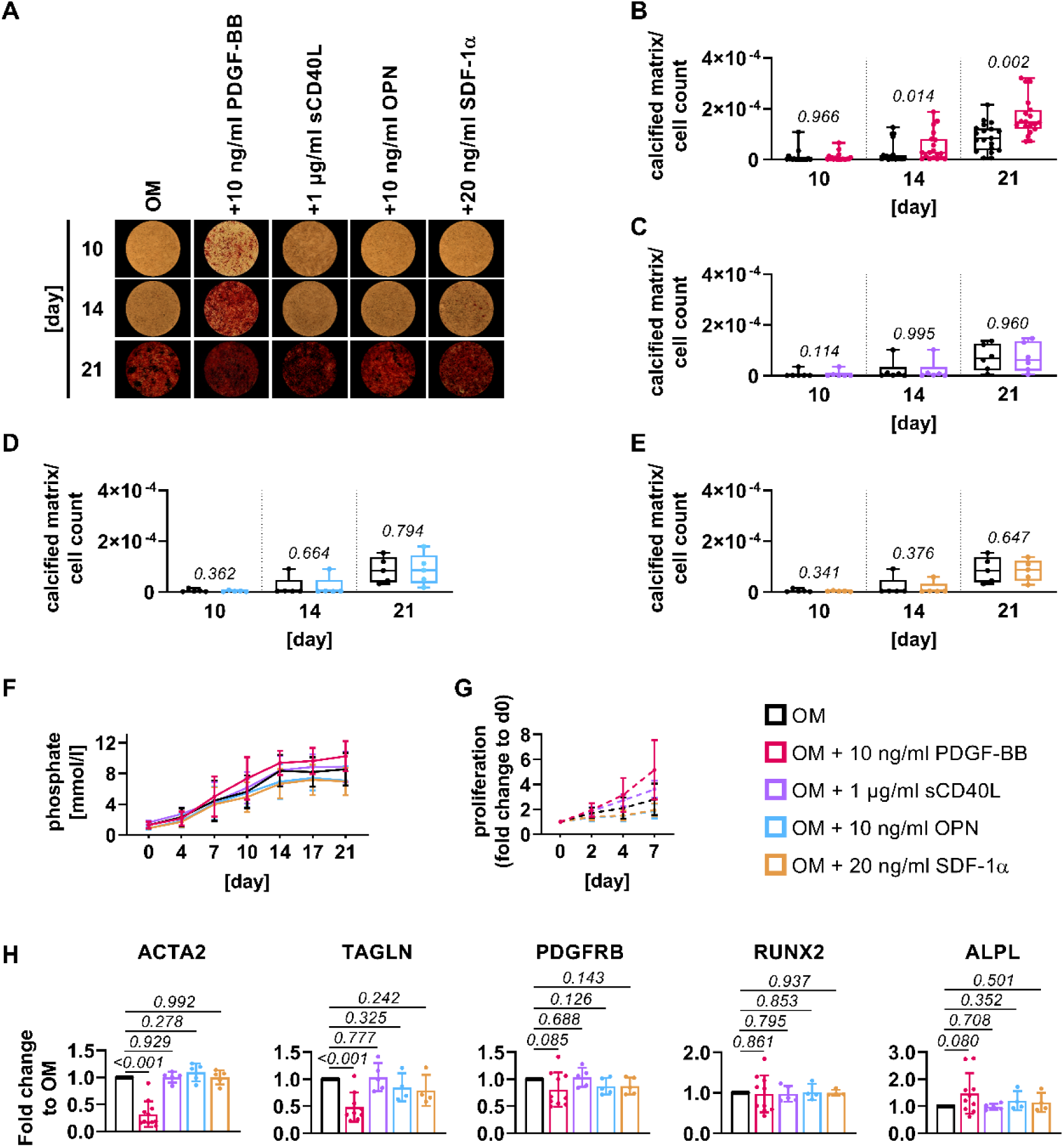
PDGF-BB increases SMC calcification. SMC undergoing osteogenic differentiation (OM - osteogenic medium) in an *in vitro* model of SMC calcification were stimulated with recombinant PDGF-BB, sCD40L, OPN, or SDF-1α. Deposited calcified matrix was stained with alizarin red. **(A**) Representative images of the deposited calcified matrix. **(B-E)** Quantified staining, normalized to cell count. A paired, two-tailed t-test was used to assess significant differences (p<0.05) vs. OM without supplements for each time point. **(F)** Phosphate levels in the supernatant during osteogenic differentiation with the respective recombinant factors. Statistical analysis at each time point was performed using a mixed-effects model with Geisser-Greenhouse correction and Dunnett’s post-test vs. OM without supplements (p<0.05); no significant differences were detected. **(G)** SMC proliferation in expansion medium (EM; dashed line) after stimulation with the respective recombinant factors. Differences in proliferation were assessed at each time point using two-way ANOVA with Geisser-Greenhouse correction and Dunnett’s post-test vs. EM without supplements (p<0.05); no significant differences were detected. **(H)** Gene expression analysis of SMC after four days of osteogenic differentiation with the respective recombinant factors. Paired, two-tailed t-tests vs. OM without supplements were used to assess significant differences (p<0.05) across independent experiments.

Consistently and in line with previous reports, the increased calcification induced by PDGF-BB or its combinations coincided with markedly increased SMC proliferation (Fig. 4 G, Extended Data Fig. 4 J) and reduced expression of SMC markers ACTA2, TAGLN, and PDGFRB, while ALPL expression increased compared to OM control (Fig. 4 H, Extended Data Fig. 4 J, Extended Data Fig. 5). In contrast, sCD40L, OPN, or SDF-1α did not affect SMC proliferation or SMC and osteogenic marker expression. Thus, PDGF-BB emerged as a potential mediator that promotes an osteogenic switch in SMC and links the systemic osteoporotic milieu with SMC calcification.

### Inhibition of PDGF-BB or its downstream signaling reduces SMC calcification induced by osteoporotic sera

To investigate causality between PDGF-BB in vascular calcification in osteoporosis, we blocked PDGF-BB activity or its downstream signaling in sex-mixed OPO^+^ and OPO^-^ sera using either a neutralizing PDGF antibody (αPDGF) or the polyphenol chebulinic acid (CheA) that was previously shown to suppress PDGFRβ phosphorylation and siganling.^33^ Neutralization of PDGF-BB with αPDGF reduced SMC calcification in cultures stimulated with OPO sera up to day 14 (Fig. 5 A, B).

**Fig. 5:**
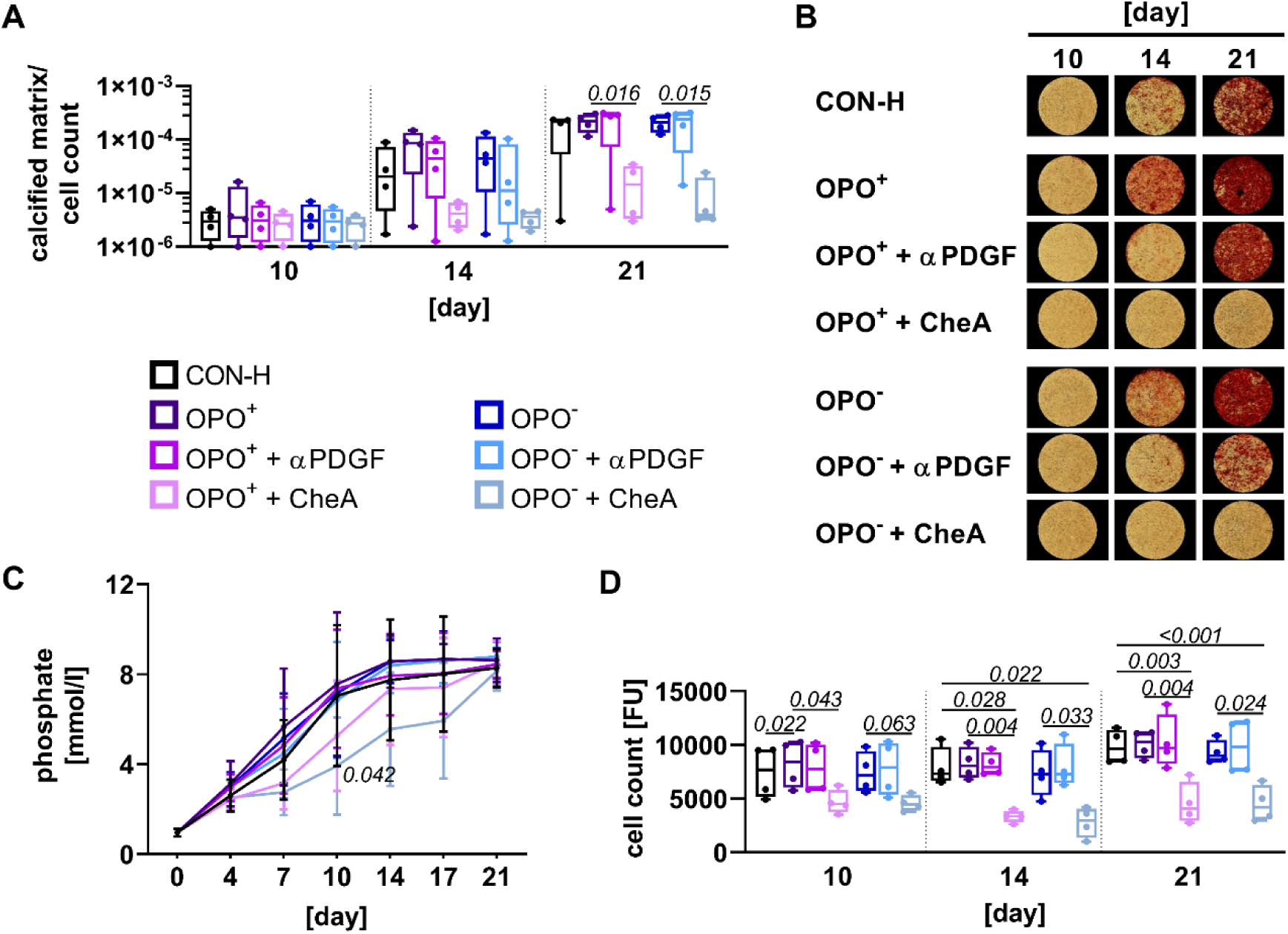
Inhibition of PDGF-BB or its downstream signaling reduces increased SMC calcification induced by osteoporotic sera. **(A)** SMC undergoing osteogenic differentiation were stimulated with sex-mixed sera pools of non-osteoporotic healthy older controls (CON-H) or osteoporotic frail individuals with (OPO^+^) or without confounders of vascular calcification (OPO^-^). PDGF-BB or its downstream signaling was inhibited in the respective sera pools using either 25 µg/ml neutralizing polyclonal PDGF antibody (αPDGF) or 50 µM chebulinic acid (CheA). Deposited calcified matrix was stained with alizarin red, quantified, and normalized to the cell count. A one-way ANOVA with Geisser-Greenhouse correction and Dunnett’s post-test was used to assess significant differences vs. CON-H and the respective osteoporotic group without inhibitor for each time point. **(B)** Representative images of deposited calcified matrix stained with alizarin red. **(C)** Phosphate levels determined in the supernatant during osteogenic differentiation. A two-way ANOVA with Geisser-Greenhouse correction and Dunnett’s post-test was used to assess significant differences vs. CON-H for each time point. **(D)** Cell counts determined by Hoechst staining after stimulation with the respective sera pools supplemented with αPDGF or CheA. A one-way ANOVA with Geisser-Greenhouse correction and Dunnett’s post-test was used to assess significant differences vs. CON-H and the respective osteoporotic group without inhibitor for each time point. Only significant differences (p<0.05) and distinct trends with p<0.1 are shown.

However, blocking PDGF-BB signaling with CheA prevented SMC calcification entirely, reduced phosphate levels even below levels observed in cultures treated with CON-H sera (Fig. 5 C), and halted SMC proliferation over the course of the differentiation (Fig. 5 D). Consistently, ERK1/2 phosphorylation, being critical for osteogenic differentiation, was reduced to baseline levels in human coronary artery SMC from a healthy donor after four days of osteogenic differentiation with PDGF-BB in the presence of CheA (Extended Data Fig. 6). Together, these findings emphasize the role of PDGF-BB as a central mediator of vascular calcification in osteoporotic individuals and suggest its signaling axis as a potential therapeutic target.

## Discussion

In this exploratory human study, we combined retrospective *ex vivo* analyses with mechanistic *in vitro* experiments to investigate the association between osteoporosis and vascular calcification. We identified a systemic osteoporotic milieu characterized by increased eosinophil, neutrophil, and platelet counts, elevated sCD40L and PDGF-BB levels, and altered SDF-1 and OPN levels. This milieu was found to correlate with markers of bone turnover and arterial stiffening, indicating that bone loss is associated with systemic immune activation, as well as local vascular alterations. Notably, we provide comprehensive human evidence that osteoporotic sera increase SMC calcification *in vitro*, driven by PDGF-BB as a direct mediator. Inhibition of PDGF-BB downstream signaling effectively prevents this process. These findings expand the current understanding of the pathological bone-vascular cross-talk in humans and substantiate PDGF-BB as one causal mediator and a promising therapeutic target.

Our systematic stratification revealed an immune cell signature in individuals with low BMD, characterized by increased granulocyte and platelet counts, as well as higher basophil-to-lymphocyte ratios, eosinophil-to-lymphocyte ratios (ELR), and neutrophil-to-lymphocyte ratios (NLR), and a higher systemic inflammation index (SII). These findings are consistent with previous reports showing increased neutrophil-to-lymphocyte ratios^34–36^ and systemic inflammation indices^31^ in postmenopausal women with low BMD and osteoporotic individuals of both sexes, as well as a negative association between the platelet-to-lymphocyte ratio and lumbar spine BMD in males.^37^ In our study, a similar male bias was observed for increased eosinophil and neutrophil counts, whereas platelet counts were increased in both sexes.

Although our data are limited in sample size and statistical power for detailed subgroup analyses, the consistent associations suggest that disrupted bone homeostasis and bone marrow activation alter hematopoiesis in both sexes,^19,38–40^ with direct vascular consequences. Earlier work has linked low BMD and increased granulocyte counts to clonal hematopoiesis of indeterminate potential (CHIP),^41^ a condition that accelerates atherosclerotic processes through clonal expansion of hematopoietic stem cells with somatic mutations.^42^ In particular high neutrophil counts have been associated with atherosclerotic cardiovascular disease.^43^ Mechanistic studies showed that neutrophil interactions with activated platelets can induce neutrophil extracellular trap formation, thereby promoting SMC death and plaque instability.^44^ In addition, cationic proteins released from eosinophils^45^ and platelet-derived signals^46^ may promote vascular remodeling and calcification.

Considering the more rapid loss of trabecular bone after menopause,^47^ longitudinal studies are needed to determine how granulocyte and platelet counts, as well as their activation status, change with progressive loss of BMD in both sexes.

Beyond the cellular signature, we observed two distinct molecular signatures in the participant-derived sera: a frailty-associated, low-grade pro-inflammatory serum milieu and an osteoporotic milieu characterized mainly by platelet-associated mediators. The latter was marked by increased sCD40L levels that closely correlate with increased platelet counts and PDGF-BB levels. This finding aligns with previous reports demonstrating that CD40L is cleaved mainly from platelets upon their activation,^48^ a process that coincides with the release of α-granule contents, including PDGF-BB.^48,49^ Accordingly, in addition to the documented release from preosteoclasts,^9,10^ our data suggest platelets as an additional source of PDGF-BB in the pathological bone-vascular cross-talk. Despite only partially available cardiovascular assessments in our retrospective study, correlations of sCD40L and PDGF-BB with IMT and PWV indicate that respective systemic alterations are also associated with arterial stiffness in humans, substantiating the previously reported link between IMT and PWV with BMD.^50^ Notably, a recent study also associated sCD40L, the CD40 receptor, and PDGF-BB with total carotid plaque area in individuals with high cardiovascular risk.^51^

The mechanistic role of PDGF-BB is further supported by our *in vitro* experiments: similar to sera pools from osteoporotic and osteopenic individuals, recombinant PDGF-BB significantly increased SMC calcification, while SMC marker expression was reduced. Neutralization of PDGF-BB or inhibition of its downstream signaling in human SMC cultures with osteoporotic sera pools reduced or suppressed this effect. Especially, inhibition of PDGF-BB downstream signaling using chebulinic acid broadly prevented SMC calcification, likely due to a reduced PDGF-BB-driven ERK1/2 phosphorylation^33^ and respective downstream regulatory effects on pro-osteogenic RUNX2 signaling.^11^ Consistent with findings in male mice,^9,10^ our data establish PDGF-BB as a central mediator linking bone loss with arterial stiffness and vascular calcification in humans.

Other altered serum factors, sCD40L, OPN, SDF-1, showed no effect on SMC calcification, but might contribute to vascular calcification indirectly by exerting effects on endothelial cells, macrophages,^48,52^ or other immune cells that cannot be studied in our current *in vitro* model. For example, it was demonstrated that the SDF-1/ CXCR4 axis maintains arterial integrity,^53^ while OPN seems to have a dualistic role as an inhibitor or enabler of mineralization depending on its phosphorylation status.^54^ An increase in serum OPN levels in postmenopausal osteoporotic women was reported previously as well,^55^ whereas positive correlations with IMT and PWV in our study establish a link to the vasculature. These previously described secondary effects together with the observed associations between bone turnover, systemic immune alterations, and arterial stiffening emphasize that vascular calcification in osteoporotic individuals might not be driven by a simple cause-and-effect relationship.

Our analyses rather indicate that the pathological bone-vascular cross-talk needs to be conceptualized as a complex adaptive system (CAS), with simultaneous interactions and potentially dynamic, interdependent cause-and-effect relationships between its components and nested subsystems as outlined in Fig. 6. Unaware of the systems behavior as a whole, these physically distant subsystems and individual components react to signals in their local environment and adapt to altered signaling cues in tissue-specific self-organizing processes.^26,27^ Thereby, some components and subsystems function as a CAS themselves,^26,27^ for example through the diverse functions of granulocytes and platelets, and the demonstrated adaptive behavior of calcifying SMC. This point of view explains how even minor, local changes in bone homeostasis, such as increased granulopoiesis and PDGF-BB released from platelets and preosteoclasts,^10^ can spread through the circulation, and manifest as pathological calcification in the arteries.^26,27,56^ Here, CAS theory highlights the role of dynamic, nonlinear interactions,^26,27^ which also seem to be of relevance in the pathological bone-vascular cross-talk as indicated by the multiple interactions between the components of the osteoporotic milieu. Accordingly, the systems behavior and the development of vascular calcifications in response to the osteoporosis-associated alterations emerges as the product of these interdependent interactions,^24^ in the attempt to find a new degree of order.^26,27,57^ The identification of PDGF-BB as a central mediator in this broader adaptive network, provides a potential molecular entry point to therapeutically shift the system towards healthier outcomes. Both targeted intervention strategies that benefit healthier aging and a comprehensive understanding of osteoporosis-associated changes over time require longitudinal study designs.

**Fig. 6:**
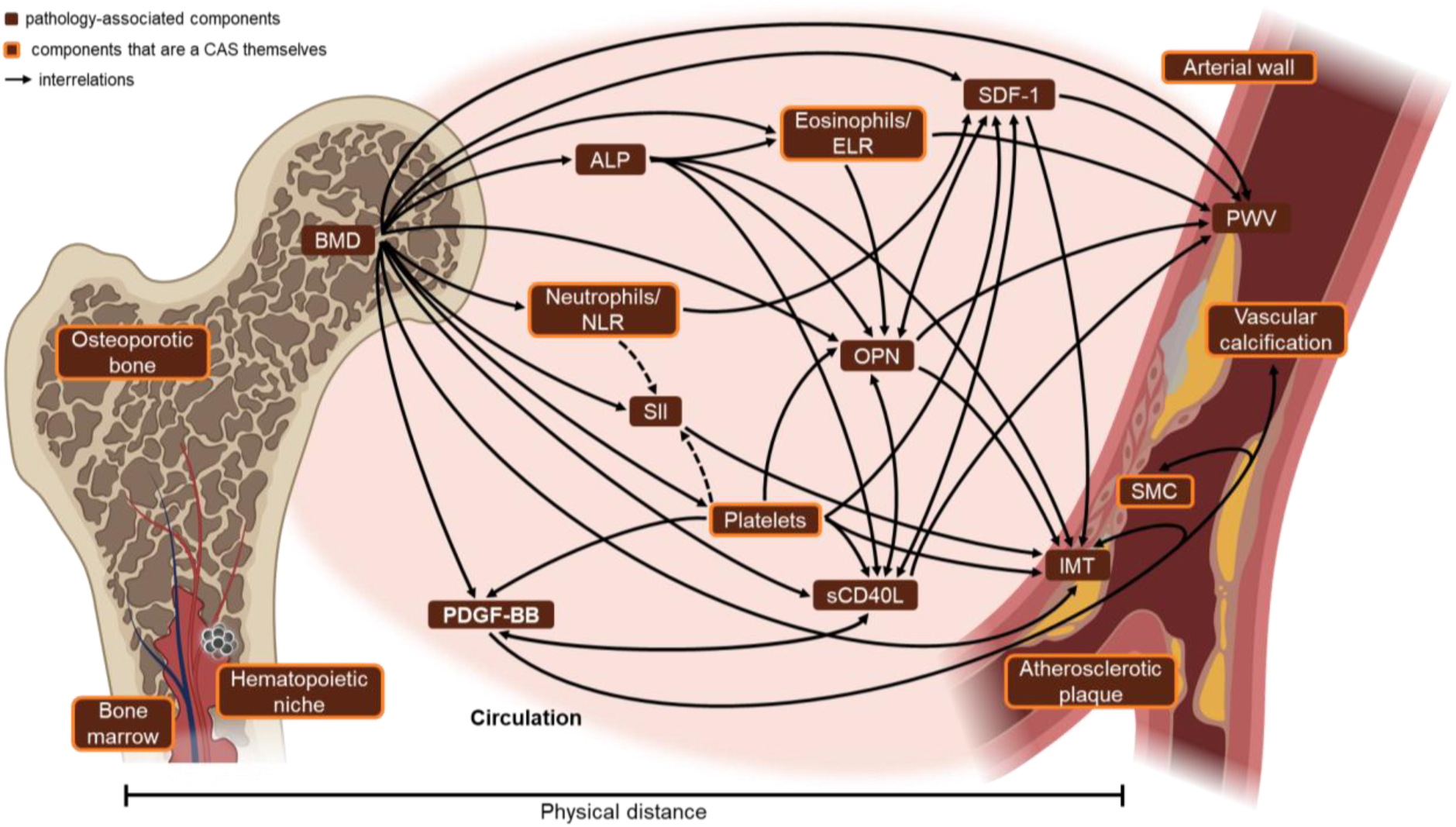
The pathological bone-vascular cross-talk evolves as a complex adaptive system. Interdependent associations of diverse components of the complex adaptive system based on *ex vivo*, *in vitro*, and correlation analyses. ALP - alkaline phosphatase, BMD - bone mineral density, ELR - eosinophil-to-lymphocyte ratio, IMT - intima-media thickness, NLR - neutrophil-to-lymphocyte ratio, PWV - pulse wave velocity, SII - systemic inflammation index (NLR × platelet count), SMC - smooth muscle cells.

## Methods

### Sample acquisition and study criteria

Data and sample collection from human participants was conducted in accordance with ICH-GCP E6 and the Declaration of Helsinki after approval by the Institutional Ethics Committees of the Medizinische Fakultät, Ruhr-Universität Bochum (OsteoSys study, approval 16-5714, 07.07.2016, amendment approved 24.01.2017; recruitment from February 2017 to October 2019) and the Charité - Universitätsmedizin Berlin (Frailty study, approval EA1/208/15, 6.06.2017, recruitment from March 2016 to March 2018 and study „Einfluss des Serums auf das regenerative Potential von mesenchymalen Stammzellen (MSC) *in vitro*“, approval EA2/101/10, 21.10.2010, recruitment from March 2018 to March 2019). Written informed consent was obtained from all participants. The OsteoSys study and the study “Einfluss des Serums auf das regenerative Potential von mesenchymalen Stammzellen (MSC) *in vitro*” collected data only at the time of enrollment, with no follow-up data available. Accordingly, from the Frailty study, only data and samples obtained at study entry were used.

At the Ruhr-Universität Bochum center, geriatric participants with primary osteopenia or osteoporosis, including potentially osteoporosis-related fractures, as well as individuals with normal BMD, were recruited from the Marien and St. Anna Hospital and the Rheumatology Center Ruhrgebiet (Herne, Germany) for the OsteoSys study. Participants with secondary osteoporosis, persistent hyperthyroidism, renal osteopathy, aromatase inhibitor or hormone-ablative antiandrogenic therapy, systemic mastocytosis, active chronic inflammatory bowel disease, type 1 diabetes, or a body mass index (BMI) <17.5 kg/cm² were excluded. At the Berlin center, participants for the Frailty study were recruited from retirement homes. Eligible individuals were ≥65 years, legally competent, and able to walk 70 meters unaided. Individuals with major medical conditions such as Parkinson’s disease, neuromuscular or musculoskeletal disorders, cardiovascular diseases, severe lung, kidney, or liver diseases, chronic viral infections (e.g. HIV or hepatitis), current or recent cancer, or autoimmune diseases such as type 1 diabetes or multiple sclerosis were excluded from the study. For the study „Einfluss des Serums auf das regenerative Potential von mesenchymalen Stammzellen (MSC) *in vitro*“, participants aged 18 to 80 years without known severe or current diseases (cancer; infections such as hepatitis, HIV, or tuberculosis; diabetes; cardiovascular, bone, lung, kidney, or liver diseases; psychiatric disorders such as major depression) were recruited.

Pregnant individuals and drug abusers were excluded. Blood and sera samples, together with available diagnostic data from all centers were used for the present study.

### Diagnostic assessment of osteoporosis, frailty, and risk factors for cardiovascular disease

Diagnostic assessment, including physical and laboratory examinations, was conducted according to the respective study protocols. At the Ruhr-Universität Bochum center, BMD was assessed by dual-energy X-ray absorptiometry (DXA) using a Lunar Prodigy Advance (GE Medical Systems, USA). Measurements were performed in the supine position at the lumbar spine and both hips (total femur and femoral head). The lowest T score of all regions was used for diagnosis according to WHO criteria: T score ≤ −2.5, osteoporosis; T score between −1 and −2.5, osteopenia; T score > −1, non-osteoporotic. Using questionnaires and medical records, previous diagnoses of osteoporosis and respective treatments, pre-existing orthopedic conditions (fractures within the last five years or fractures with long-term consequences, joint replacement, meniscal surgeries, osteoarthritis, tendon ruptures, sarcopenia), the number of falls in the last year, and potential frailty-related difficulties in climbing one level of stairs were assessed. As part of the frailty assessment, HGS of the dominant or unaffected hand was measured using a Jamar hydraulic hand dynamometer (Lafayette Instrument Company, USA). HGS measurements were performed three times, and the maximum value was recorded.

Pre-existing strokes, as well as cardiovascular events and diseases, were documented by questionnaire. In addition, PWV and IMT were determined. PWV was measured in supine position after 5 min of rest using a SphygmoCor^®^ XCEL (AtCor Medical, Australia, software version 1.2). Brachial blood pressure, obtained by the standard cuff oscillometric method, was used for calibration, and the median PWV was calculated from three consecutive measurements. IMT was assessed by carotid ultrasound imaging (Aloka/Hitachi, Prosound alpha 6, Dormed Hellas SA, Greece) with a 5 - 13 MHz high-frequency linear-array transducer. Longitudinal B-mode images of both common carotid arteries were recorded at end-diastole and mean and maximum thickness were determined at the continuous region 10 mm proximal to the bulb origin at the far wall using an auto-tracing system. The average of both sides was calculated.

Clinical assessment further included documentation of age, height, weight, current and previous smoking status, blood pressure, as well as blood levels of calcium, phosphate, ALP, (25)-Hydroxy-Vitamin D, low- and high-density lipoprotein cholesterol (LDL-C and HDL-C), and platelet counts. Pre-existing diagnoses and medication for dementia, diabetes mellitus, hyperlipidemia, hyperuricemia, renal insufficiency, and rheumatoid arthritis were also documented by questionnaire.

Within the Frailty study at the Charité - Universitätsmedizin Berlin, HGS of the dominant hand was measured twice using a Jamar Plus digital hand dynamometer, with the maximum value recorded. Age, height, weight, as well as current and previous smoking status were documented and platelet counts were determined by Labor Berlin - Charité Vivantes GmbH (Germany). Pre-existing diagnoses and medication for dementia, diabetes mellitus, hypertension, stroke, cardiovascular events and diseases, rheumatoid arthritis, orthopedic conditions (as outlined above), osteochondrosis, and disc prolapses, as well as the number of falls in the last year, were recorded by questionnaire. Stair-climbing ability was evaluated in a ten-step up-and-down test; all participants selected for this study completed the test in <10 seconds. For comparison across studies, these results were classified as “no difficulty climbing stairs”.

### Flow cytometry analysis

Peripheral blood was collected in S-Monovette K3 EDTA blood collection tubes (Sarstedt). For staining, 25 µl whole blood was mixed with 25 µl staining solution (Dulbecco’s Phosphate Buffered Saline (PBS; Life Technologies) containing antibodies listed in Supplementary Table 1) and incubated for 10 min at room temperature in the dark. All antibodies were titrated to determine optimal concentrations. Subsequently, erythrocytes were lysed adding 450 µl of VersaLyse (Beckman-Coulter, USA) and incubated for 15 min at room temperature in the dark. Samples were then centrifuged at 400 x g for 10 min and resuspended in 200 µl PBS. Immediately before acquisition, 1 µg/ml propidium iodide (Thermo Fisher, USA) was added to distinguish living from dead cells. Acquisition was performed on a CytoFlex flow cytometer (Beckman Coulter). Instrument performance was verified daily using CytoFlex Daily QC Fluorospheres (Beckman Coulter). No modification of compensation matrices was required throughout the study. Data were analyzed using FlowJo V10.6.1 (BD Bioscience, USA). Determined subset counts were used for calculation of ratios to overall lymphocyte counts. No platelet-to-lymphocyte ratio was not calculated, as platelet counts were determined separately as part of the clinical assessment, independent of flow cytometry.

### Serum screening

Sera were screened using bead-based multiplex assays (LEGENDplex^TM^, BioLegend^®^, USA) according to the manufacturer’s instructions. Reagent and sample volumes were reduced to 10 μl following correspondence with the manufacturer. The pre-defined Inflammation Panel 1 and Proinflammatory Chemokine Panel 1, as well as a customized panel including APRIL, BAFF, sCD40L, CTLA-4, IL-4, IL-9, OPG, OPN, PDGF-BB, RANKL, and SDF-1 (CXCL12) were used. Data were acquired on a CytoFlex S or LX flow cytometer (both Beckman Coulter) and analyzed using the LEGENDplex™ Data Analysis Software provided by the manufacturer. Expression levels below the detection range of the assay were set to the respective lower detection limit.

### SMC culture and osteogenic differentiation

For the mechanistic *in vitro* studies, a previously established model of vascular calcification was used that incorporates a broad and clinically relevant variance of human SMC *from different locations and donors.*^32^ *SMC isolated from* total endarterectomy samples of calcified arteries and aneurysms of the femoral and iliofemoral arteries, as well as a shunt vein revision were used for this study (four female donors: 68 to 95 years, two male donors: 66 and 76 years; tissue samples from Marien Hospital Herne, Germany; ethical approval 16-5714, 07.07.2016, amendment approved 24.01.2017). Cells were expanded in Dulbecco’s Modified Eagle Medium low glucose (DMEM, D5546; Sigma-Aldrich, Germany) supplemented with 1% GlutaMAX^TM^ (Life Technologies, Netherlands), 1% penicillin/streptomycin (Biochrom, Germany), and 10% fetal calf serum (FCS; Biochrom). In addition, three commercially available healthy coronary artery SMC donors (one female donor: 34 years; two male donors: 20 and 46 years; Lifeline^®^ Cell Technology, USA) were included after gradual transfer from VascuLife SMC Medium (Lifeline^®^ Cell Technology) to DMEM during expansion. All cells were tested negative for mycoplasms.

SMC at passage four were seeded either subconfluently into 96 well plates (Falcon^®^, Corning, USA) for proliferation assay or as a near-confluent cell layers for osteogenic differentiation using DMEM expansion medium with FCS. Five replicates per condition were seeded. For gene expression analysis, 150,000 SMC/well were seeded in duplicate into 6 well plates (Falcon^®^). Osteogenic differentiation and proliferative effects were investigated according to an updated version of a previously described protocol.^32^ Osteogenic differentiation was induced 24 h post-seeding using DMEM supplemented with 1% GlutaMAX^TM^, 1% penicillin/streptomycin, 50 μM L-Ascorbic acid 2-phosphate sesquimagnesium salt hydrate, 10 mM β-Glycerophosphate disodium salt hydrate, and 0.1 μM water soluble dexamethasone (the last three all Sigma-Aldrich). To study the effect of sera from non-osteoporotic, osteopenic, and osteoporotic individuals, osteogenic medium was supplemented with 10% participant-derived pooled sera. Sex-mixed pools of equal volumes per individual were prepared to represent general disease states. Additionally, female- and male-specific pools were generated from the CON-H and OPO^+^ groups to assess sex differences. Sera pools were aliquoted, stored at −80 °C, and freshly thawed at each media exchange.

To investigate potential serum mediators identified in the *ex vivo* screening, osteogenic medium containing 10% heat-inactivated human AB serum (hAB; Sigma-Aldrich) was supplemented with recombinant human sCD40L (1 µg/ml; PeproTech, USA), PDGF-BB (10 ng/ml; carrier-free; BioLegend^®^), OPN (10 ng/ml; PeproTech), SDF-1α (20 ng/ml; PeproTech), or their combinations. Identically supplemented DMEM expansion medium with 10% hAB was used for proliferation assays.

The role of PDGF-BB was further assessed using osteogenic medium containing sex-mixed participant-derived sera pools supplemented with either 25 µg/ml neutralizing polyclonal human PDGF antibody (αPDGF; Cat# AB-20, RRID:AB_354273; R&D Systems^®^) dissolved in PBS, or 50 µM chebulinic acid (CheA; Cat# SMB00271, Sigma-Aldrich or Cat# A16208, AdooQ^®^ BioScience, USA), dissolved in ddH_2_O and briefly heated up to 70 °C to prepare a 1 mM stock. Media with PDGF-BB inhibitors and corresponding controls were pre-incubated for 1 h at 37 °C before being added to the *in vitro* cultures. All recombinant factors and inhibitors were reconstituted according to the manufacturer’s instructions, aliquoted, and stored at −20 °C until use. Osteogenic and proliferation media were freshly prepared at each medium exchange and media were exchanged every three to four days. Proliferation was monitored until day ten, while osteogenic differentiation was continued until day 21.

### Determination of metabolic activity and proliferation

The metabolic activity of SMC was assessed prior to determining cell count and ALP activity. One volume of PrestoBlue Cell Viability Reagent (Life Technologies) was mixed with ten volumes of pre-warmed DMEM expansion medium with FCS, and SMC were incubated for 1 h at 37 °C and 5% CO_2_ with 64 μl/96 well of this mixture.

Absorbance was measured at 590 nm using an Infinite^®^ M 200 PRO plate reader (Tecan Group Ltd., Switzerland). Blank values were subtracted, and metabolic activity was either normalized to the cell count of each SMC donor or used for normalization of ALP activity. Data are presented as mean ± standard deviation (SD) for each SMC donor and condition.

Proliferation of SMC was assessed at baseline (day zero) and on days two, four, and seven. After determination of metabolic activity, cells were washed once with PBS, and plates were stored empty at −80 °C until analysis. Cell proliferation was determined using the CyQUANT^TM^ cell proliferation assay (Life Technologies) according to the manufacturer’s instructions. Blank values were subtracted, and cell counts were calculated for each SMC donor as mean fold change relative to day zero per condition ± SD.

### Evaluation of deposited calcified matrix

Calcified matrix deposited by osteogenically stimulated SMC was assessed by alizarin red staining after 10, 14, and 21 days of differentiation. Cells were washed with PBS and fixed with ROTI^®^Histofix 4% (Carl Roth, Germany) for 10 min at room temperature. After removing the fixative, cells were washed with PBS and covered with 75 µl fresh PBS for Hoechst background measurement at 460 nm using an Infinite® M 200 PRO plate reader. Cells were then stained with 10 μg/ml Hoechst (Sigma-Aldrich) diluted in PBS for 10 min at room temperature, washed once with PBS, and covered with 75 µl fresh PBS per well. Cell counts were determined at 460 nm using the same plate reader. For data analysis, background and blank values were subtracted, and mean cell counts ± SD were calculated per SMC donor and condition.

Subsequently, cells were washed once with ddH_2_O and stained with 75 µl 0.5% alizarin red S (Sigma-Aldrich; in ddH_2_O, pH 4.2) for 10 min at room temperature. Unbound dye was removed by three washes with ddH₂O, with additional washes performed as needed. Plates were dried, and staining of calcified matrices was documented using a REBEL REB-01 microscope (ECHO, USA) equipped with a 4x plan achromatic phase contrast objective, a 12MP CMOS Color camera, and the imaging software ECHO Pro V4.5. For quantification, alizarin red staining was eluted with 106 µl/well cetylpyridinium chloride (Sigma-Aldrich; 10% solution in ddH_2_O).

From this 100 µl of eluate were transferred to a new 96 well plate (Sarstedt), and optical density was determined at 562 nm using the Infinite^®^ M 200 PRO plate reader. Blank values of cetylpyridinium chloride were subtracted, and mean alizarin red values ± SD were calculated per SMC donor and condition.

### Determination of ALP activity and phosphate levels

Enzymatic ALP activity of SMC and phosphate levels in the culture supernatant were assessed as markers of osteogenic differentiation. ALP activity was determined on day zero, seven, and ten, after assessment of metabolic activity, by quantifying para-nitrophenyl phosphate disodium hexahydrate (*p*NPP) turnover to 4-nitrophenolate.

Buffers and substrate solution were preheated to 37 °C. Cells were washed once with 150 μl PBS, followed by 200 μl AP-buffer (100 mM NaCl, 100 mM Tris(Hydroxymethyl-)aminomethane (both Merck, Germany), 1 mM MgCl_2_ (Carl Roth), 100 ml ddH_2_O, pH adjusted to 9.0). Cells were then incubated with 100 μl *p*NPP substrate (*p*NPP solution: 1 mg/ml *p*NPP dissolved in 1 M diethanolamine (both Sigma-Aldrich), add 100 ml ddH_2_O, pH adjusted to 9.8; mix 1:1 with AP-buffer) for 10 min at 37 °C and 5% CO_2_. The reaction was terminated by addition of 100 µl 1 M sodium hydroxide solution (Sigma-Aldrich). From each well, 100 µl of supernatant were transferred to a new 96 well plate (Sarstedt), and absorbance was measured at 405 nm using an Infinite^®^ M 200 PRO plate reader. Blank values were subtracted. *p*NPP turnover was calculated using a molar extinction coefficient of 18,450 l x mol^-1^ x cm^-1^, and ALP activity was defined as nmol 4-nitrophenolate formed per min at 37 °C. ALP activity was normalized to cellular metabolic activity and is expressed as fold change relative to baseline at day zero per SMC donor and condition ± SD.

Phosphate levels were determined in cell culture supernatants collected from SMC undergoing osteogenic differentiation at each medium exchange and stored at −80 °C until analysis. Phosphate was measured using the Phosphate Assay Kit (Abcam, UK) according to the manufacturer’s instructions, with absorbance read at 650 nm on an Infinite® M 200 PRO plate reader. Blank values were subtracted, and phosphate concentrations were calculated as mean ± SD per SMC donor and condition.

### Gene expression analysis

For gene expression analysis, cells were placed on ice, washed twice with cold PBS (4 °C), and lysed with TRI Reagent^®^ (Zymo Research, USA). Lysates from duplicate wells per condition were pooled and snap-frozen in liquid nitrogen. RNA was isolated using the Direct-zol^TM^ RNA Mini- or Microprep Kit (Zymo Research, USA) according to the manufacturer’s instructions. RNA concentration was determined with a NanoDrop ND-1000 spectrophotometer (PeqLab, Germany), and 500 ng RNA were reverse transcribed into cDNA using the iScript cDNA Synthesis Kit (Bio-Rad Laboratories GmbH, Germany). Quantitative real-time PCR (qRT-PCR) was performed with 5 ng of cDNA per reaction using SYBR^®^ Green PCR master mix (Roche, Switzerland) and primers designed with NCBI Primer-BLAST (Supplementary Table 2). Reactions were run on a LightCycler^®^480 II (Roche). Gene expression was analyzed using the ΔΔCt method with normalization to the housekeeping gene *RPL13A* and correction for primer efficiency. Results were expressed as fold change relative to the osteogenic medium control of each donor.

### Analysis of phosphorylation profiles

Kinase phosphorylation in human coronary artery SMC from one healthy donor (male, age: 46 years) was analyzed using the Proteome Profiler Human Phospho-Kinase Array Kit (R&D Systems®, USA). During expansion, SMC were gradually transferred from VascuLife SMC Medium to DMEM expansion medium with FCS. For differentiation, 150.000 SMC were seeded per 6 well (Falcon®, Corning) with DMEM containing 1% GlutaMAX^TM^, 1% penicillin/streptomycin, and 10% heat-inactivated hAB serum. After 24 h, the medium was replaced with DMEM containing 1% GlutaMAX^TM^, 1% penicillin/streptomycin, and 0.1% heat-inactivated hAB serum for a 24 h starvation period (baseline). Phosphorylation levels were assessed at baseline and after 15 min and four days of differentiation with osteogenic medium alone or supplemented with 10 ng/ml PDGF-BB and either 25 µg/ml neutralizing polyclonal PDGF antibody or 50 µM chebulinic acid. Lysates from three replicate wells per condition were pooled and stored at −80 °C. For analysis, lysates were thawed on ice and protein concentration determined using the Quick Start^TM^ Bradford Dye Reagent (Bio-Rad Laboratories). Absorbance was measured at 595 nm with an Infinite® M 200 PRO plate reader. Protein concentrations were calculated from a bovine serum albumin (Sigma-Aldrich) standard curve in lysis buffer, with blanks subtracted. For each array, 152 µg protein was applied. Membranes were imaged with a ChemiDoc™ MP Imaging System, and spot intensities quantified using Image Lab V6.0 software (both Bio-Rad Laboratories).

### Statistical analysis and correlation

This study was designed as a retrospective, exploratory analyses, accordingly, sample sizes were not determined by a priori power calculations. Differences in group sizes were addressed by reporting percentages or means, and sex-specific analyses were performed where possible.

Categorical diagnoses and discrete counts, e.g. comorbidities and number of falls were transformed into binary variables, ≥1 diagnosis or fall was coded as “yes” and none as “no”, to enable comparability across groups. Exploratory statistical analyses were conducted in line with common practice to identify trends, with results interpreted in the context of limited power given the small group sizes, high biological variance, limited statistical power, and potential violations of test assumptions.

The choice of statistical tests reflected data type, group size, and variance, and all applied tests are specified in figure and table legends. Briefly, normality was assessed using Anderson-Darling, D’Agostino-Pearson, Shapiro-Wilk, and Kolmogorov-Smirnov tests. Parametic tests were applied when normal distribution was supported, otherwise non-parametric tests were used. To account for small sample sizes and unequal variance, or two-tailed t tests with Welch’s correction, Brown-Forsythe and Welch ANOVA, or Geisser-Greenhouse correction, were applied. Analyses focused primarily on comparisons with CON-H, but included comparisons to all three control groups where possible. *Ex vivo* data were assessed using Brown-Forsythe and Welch ANOVA, Kruskal-Wallis tests, or two-way ANOVA with Dunnett’s or Dunn’s post-tests. Sex differences between individuals of the same study group were assessed using Sidak post-tests. Binary variables were evaluated by Fisher’s exact test with Bonferroni correction. Associations between clinical data and *ex vivo* data were calculated using Spearman correlations, with differences between correlation coefficients tested by Fisher’s Z-test.

*In vitro* assays were analyzed according to their experimental design: unpaired two-tailed t tests with Welch’s correction for SMC calcification and ALP activity in serum-stimulated SMC cultures of independent experiments, one-way ANOVA with Dunnett’s post-test for PDGF-BB inhibitor experiments, paired two-tailed t tests for SMC calcification and gene expression in SMC after stimulation with recombinant factors, and one- or two-way ANOVA with Geisser-Greenhouse correction or mixed-effects models with Dunnett’s post-test for phosphate levels and proliferation data. A p-value <0.05 was considered statistically significant and trends (0.05 ≥ p ≤ 0.1) were reported where relevant.

All calculations, statistical analyses, and visualizations were performed using Microsoft Excel, GraphPad Prism v8.4.3, and R Studio 2024.12.1. Heat maps based on Z-scores were generated with ClustVis. The overview of the interdependent coherences was created using biorender.com and Microsoft PowerPoint.

### Reporting summary

Further information on the research design is available in the Nature Portfolio Reporting Summary linked to this article.

## Data availability

Data underlying this study are available to other researchers upon reasonable request.

## Supporting information

STROBE Checklist

## Acknowledgments

The authors are grateful to Anastazja Andrzejewska and Eva Kohut for their valuable support with sample and data collection and Sviatlana Kaliszczyk for her technical assistance. We thank Dr. rer. nat. Levent Akyüz for his support in the sera analysis and Dr. rer. nat. Julia Löffler for reviewing the manuscript. This work was supported by grants of the Bundesministerium für Bildung und Forschung (BMBF), the Deutsche Forschungsgemeinschaft (DFG), the Europäischer Fonds für regionale Entwicklung (EFRE), and the Einstein Foundation through funding of the Berlin Institute of Health Center for Regenerative Therapies (BCRT; BMBF: FKZ 1315848A), the Berlin School for Regenerative Therapies (BSRT; DFG: GSC203), and the Einstein Center for Regenerative Therapies (EZ-2016-289), as well as by project funding from BMBF DIMEOS (01EC1402B), BMBF e:KID (01ZX1612A), DFG FG2165 (GE2512/2-2), DFG CRC1444, EFRE OsteoSys (EFRE-0800411 to 0800414, EFRE-0800427), and by the research grant FORUM of the Ruhr-Universität Bochum, Germany.

## Ethics declaration

Bjoern Buehring has no relevant COI for this paper. He has received research support, consulting fees and/or honoraria from AbbVie, Alexion, AlfaSigma, Amgen, Astra-Zeneca, Biogen, BMS, Boehringer Ingelheim, Fresenius Kabi, GE/Lunar, Janssen, Galapagos, Gilead, Hexal/Sandoz, Medimaps, MSD, Sanofi Genzyme, Theramex, and UCB in the last 5 years. Doruk Akgün received a consultant fee from Arthrex GmbH and Medacta International. All other authors declare no competing interests.

## Supplementary data

Extended Data Table 1-4

Extended Data Figure 1-6

Supplementary Figure 1

Supplementary Tables 1 and 2

STROBE Checklist

