## Extended Data Figures for "A complex osteoporotic milieu is associated with arterial stiffening and PDGF-BB-mediated calcification of human smooth muscle cells"

**Extended Data Table 1: Diagnostic data of female non-osteoporotic, osteopenic, and osteoporotic individuals.**

Data are given as n, mean ± standard deviation, overall percentages, or percentages of individuals presenting with more than one condition. Differences between study groups were analyzed using a Brown-Forsythe and Welch ANOVA with Dunnett's post-test for numerical data or Fisher's exact test with Bonferroni correction for binary variables (p<0.05 vs. CON-M, CON-H, or CON-F).

| Study groups | CON-H | CON-F | OPE | OPO | CON-M |
|---|---|---|---|---|---|
| Study center | Charité – Universitäts-medizin Berlin | Ruhr-Universität Bochum | | | Charité – Universitäts-medizin Berlin |
| **General** | | | | | |
| n female | 12 | 8 | 8 | 19 | 7 |
| Age [years] | 72.2 ± 4.8 | 65.3 ± 11.2 | 70.1 ± 12.7 | 72.8 ± 10.1 | 45.5 ± 12.9 |
| *vs. CON-M* | *p=0.009* | *p=0.048* | *p=0.017* | *p=0.008* | |
| *vs. CON-H* | | *p=0.403* | *p=0.985* | *p=0.999* | *p=0.009* |
| *vs. CON-F* | *p=0.403* | | *p=0.880* | *p=0.392* | *p=0.048* |
| BMI [kg/m²] | 24.1 ± 3.0 | 28.5 ± 6.1 | 26.6 ± 5.2 | 24.1 ± 4.1 | na |
| *vs. CON-H* | | *p=0.224* | *p=0.525* | *p>0.999* | |
| *vs. CON-F* | *p=0.224* | | *p=0.874* | *p=0.232* | |
| **Osteoporosis and frailty assessment** | | | | | |
| BMD [T score] | no known osteoporosis | 0.0 ± 0.9 | -2.0 ± 0.4 | -3.3 ± 0.6 | no known osteoporosis |
| *vs. CON-F* | | | *p<0.001* | *p<0.001* | |
| ≥ 1 Pre-existing orthopedic condition§ [%] | 63.6 | 87.5 | 87.5 | 47.4 | na |
| *vs. CON-H* | | *p>0.999* | *p>0.999* | *p>0.999* | |
| *vs. CON-F* | *p>0.999* | | *p>0.999* | *p=0.270* | |
| | (n = 11/12) | (n = 8/8) | (n = 8/8) | (n = 19/19) | |
| ≥ 1 Osteochondrosis & Disc prolapse§ [%] | 10.0 | 50.0 | 37.5 | 5.3 | na |
| *vs. CON-H* | | *p=0.353* | *p=0.823* | *p>0.999* | |
| *vs. CON-F* | *p=0.353* | | *p>0.999* | *p=0.052* | |
| | (n = 10/12) | (n = 8/8) | (n = 8/8) | (n = 19/19) | |
| Rheumatoid arthritis [%] | 0.0 | 25.0 | 12.5 | 10.5 | na |
| *vs. CON-H* | | *p=0.442* | *p>0.999* | *p>0.999* | |
| *vs. CON-F* | *p=0.442* | | *p>0.999* | *p>0.999* | |
| ≥ 1 Fall within last year§ [%] | 18.2 | 12.5 | 50.0 | 42.1 | na |
| *vs. CON-H* | | *p>0.999* | *p=0.957* | *p=0.739* | |
| *vs. CON-F* | *p>0.999* | | *p=0.846* | *p=0.603* | |
| | (n = 11/12) | (n = 8/8) | (n = 8/8) | (n = 19/19) | |
| Difficulty climbing stairs [%] | 0.0 | 87.5 | 75.0 | 57.9 | na |
| *vs. CON-H* | | *p<0.001* | *p=0.002* | *p=0.004* | |
| *vs. CON-F* | *p<0.001* | | *p>0.999* | *p=0.603* | |
| ≥ 1 ∑ numerical frailty assessment [%] | 83.3 | 100.0 | 100.0 | 94.7 | na |
| *vs. CON-H* | | *p>0.999* | *p>0.999* | *p>0.999* | |
| *vs. CON-F* | *p>0.999* | | *p>0.999* | *p>0.999* | |
| HGS/BMI | 1.115 ± 0.30 | 1.089 ± 0.49 | 1.002 ± 0.46 | 1.037 ± 0.39 | na |
| *vs. CON-H* | | *p=0.999* | *p=0.898* | *p=0.890* | |
| *vs. CON-F* | *p=0.999* | | *p=0.975* | *p=0.990* | |

**Continuation Extended Data Table 1**

| Study groups | CON-H | CON-F | OPE | OPO | CON-M |
|---|---|---|---|---|---|
| **Clinical laboratory diagnostics** | | | | | |
| Calcium [mmol/l] | na | 2.35 ± 0.16 | 2.28 ± 0.12 | 2.27 ± 0.13 | na |
| *vs. CON-F* | | | *p=0.483* | *p=0.387* | |
| Phosphate§ [mg/dl] | na | 3.61 ± 0.52 | 3.78 ± 0.82 | 3.55 ± 0.53 | na |
| *vs. CON-F* | | | *p=0.869* | *p=0.951* | |
| | | (n = 8/8) | (n = 8/8) | (n = 18/19) | |
| ALP§ [IU/l] | na | 75.4 ± 18.7 | 97.8 ± 53.3 | 95.4 ± 38.4 | na |
| *vs. CON-F* | | | *p=0.485* | *p=0.163* | |
| | | (n = 8/8) | (n = 8/8) | (n = 18/19) | |
| VitD§ [ng/ml] | na | 30.0 ± 19.2 | 11.4 ± 5.7 | 20.9 ± 11.0 | na |
| *vs. CON-F* | | | *p=0.057* | *p=0.417* | |
| | | (n = 8/8) | (n = 8/8) | (n = 18/19) | |
| LDL-C§ [mg/dl] | na | 138.8 ± 33.6 | 130.4 ± 39.0 | 119.4 ± 42.0 | na |
| *vs. CON-F* | | | *p=0.876* | *p=0.399* | |
| | | (n = 8/8) | (n = 8/8) | (n = 18/19) | |
| HDL-C§ [mg/dl] | na | 75.1 ± 23.3 | 57.1 ± 14.3 | 63.3 ± 18.9 | na |
| *vs. CON-F* | | | *p=0.162* | *p=0.400* | |
| | | (n = 8/8) | (n = 8/8) | (n = 18/19) | |
| **Cardiovascular assessment** | | | | | |
| IMT§ [mm] | na | 0.750 ± 0.18 | 0.814 ± 0.18 | 0.892 ± 0.24 | na |
| *vs. CON-F* | | | *p=0.871* | *p=0.257* | |
| | | (n = 8/8) | (n = 7/8) | (n = 12/19) | |
| PWV§ [m/s] | na | 4.88 ±1.41 | 7.32 ± 3.87 | 6.09 ± 1.35 | na |
| *vs. CON-F* | | | *p=0.632* | *p=0.345* | |
| | | (n = 5/8) | (n = 5/8) | (n = 7/19) | |
| pSBP [mm Hg] | na | 140.0 ± 23.3 | 137.6 ± 9.9 | 131.5 ± 13.9 | na |
| *vs. CON-F* | | | *p=0.957* | *p=0.581* | |
| pDBP [mm Hg] | na | 84.4 ± 12.7 | 86.6 ± 15.1 | 81.5 ± 10.6 | na |
| *vs. CON-F* | | | *p=0.936* | *p=0.824* | |
| **Previous cardiovascular diagnoses** | | | | | |
| Hypertension§ [%] | 30.0 | 62.5 | 75.0 | 52.6 | na |
| *vs. CON-H* | | *p>0.999* | *p=0.460* | *p>0.999* | |
| *vs. CON-F* | *p>0.999* | | *p>0.999* | *p>0.999* | |
| | (n = 10/12) | (n = 8/8) | (n = 8/8) | (n = 19/19) | |
| Stroke [%] | 0.0 | 0.0 | 0.0 | 26.3 | na |
| *vs. CON-H* | | *p>0.999* | *p>0.999* | *p=0.385* | |
| *vs. CON-F* | *p>0.999* | | *p>0.999* | *p=0.840* | |
| Coronary artery disease [%] | 8.3 | 0.0 | 0.0 | 26.3 | na |
| *vs. CON-H* | | *p>0.999* | *p>0.999* | *p>0.999* | |
| *vs. CON-F* | *p>0.999* | | *p>0.999* | *p=0.840* | |
| Cardiac infarction [%] | 8.3 | 0.0 | 0.0 | 0.0 | na |
| *vs. CON-H* | | *p>0.999* | *p>0.999* | *p>0.999* | |
| *vs. CON-F* | *p>0.999* | | *p>0.999* | *p>0.999* | |
| Cardiac insufficiency [%] | 0.0 | 0.0 | 0.0 | 10.5 | na |
| *vs. CON-H* | | *p>0.999* | *p>0.999* | *p>0.999* | |
| *vs. CON-F* | *p>0.999* | | *p>0.999* | *p>0.999* | |

**Continuation Extended Data Table 1**

| Study groups | CON-H | CON-F | OPE | OPO | CON-M |
|---|---|---|---|---|---|
| **Continuation previous cardiovascular diagnoses** | | | | | |
| Peripheral artery disease [%] | 0.0 | 0.0 | 0.0 | 5.3 | na |
| *vs. CON-H* | | *p>0.999* | *p>0.999* | *p>0.999* | |
| *vs. CON-F* | *p>0.999* | | *p>0.999* | *p>0.999* | |
| ≥ 1 ∑ previous cardio-vascular diagnoses [%] | 33.3 | 62.5 | 75.0 | 63.2 | na |
| *vs. CON-H* | | *p>0.999* | *p=0.509* | *p=0.447* | |
| *vs. CON-F* | *p>0.999* | | *p>0.999* | *p>0.999* | |
| **Assessment of secondary diagnoses and related medication** | | | | | |
| Diabetes [%] | 0.0 | 0.0 | 0.0 | 5.3 | na |
| *vs. CON-H* | | *p>0.999* | *p>0.999* | *p>0.999* | |
| *vs. CON-F* | *p>0.999* | | *p>0.999* | *p>0.999* | |
| Use of blood sugar reducing medication [%] | 0.0 | 0.0 | 0.0 | 5.3 | na |
| *vs. CON-H* | | *p>0.999* | *p>0.999* | *p>0.999* | |
| *vs. CON-F* | *p>0.999* | | *p>0.999* | *p>0.999* | |
| Use of statins [%] | 0.0 | 0.0 | 12.5 | 10.5 | na |
| *vs. CON-H* | | *p>0.999* | *p>0.999* | *p>0.999* | |
| *vs. CON-F* | *p>0.999* | | *p>0.999* | *p>0.999* | |
| Hyperlipidemia [%] | na | 0.0 | 12.5 | 10.5 | na |
| *vs. CON-F* | | | *p>0.999* | *p>0.999* | |
| Hyperuricemia§ [%] | na | 0.0 | 0.0 | 0.0 | na |
| *vs. CON-F* | | | *p>0.999* | *p>0.999* | |
| | | (n = 8/8) | (n = 7/8) | (n = 19/19) | |
| Renal insufficiency [%] | na | 0.0 | 0.0 | 15.8 | na |
| *vs. CON-F* | | | *p>0.999* | *p>0.999* | |
| Current/previous smoker§ [%] | 20.0 | 25.0 | 25.0 | 26.3 | na |
| *vs. CON-H* | | *p>0.999* | *p>0.999* | *p>0.999* | |
| *vs. CON-F* | *p>0.999* | | *p>0.999* | *p>0.999* | |
| | (n = 10/12) | (n = 8/8) | (n = 8/8) | (n = 19/19) | |

CON-H - non-osteoporotic older healthy controls, CON-F - non-osteoporotic frail controls, OPE - osteopenic frail individuals, OPO - osteoporotic frail individuals, CON-M - non-osteoporotic middle-aged controls, n - number, BMD - bone mineral density, BMI - body mass index, ALP - alkaline phosphatase, VitD - (25)-Hydroxy-Vitamin D, LDL-C - low-density lipoprotein cholesterol, HDL-C - high-density lipoprotein cholesterol, no known osteoporosis - self-declaration corresponding to past medical examinations, pSBP - peripheral systolic blood pressure, pDBP - peripheral diastolic blood pressure, IMT - intima-media thickness, PWV - pulse wave velocity, excluded - according to the exclusion criterion applied during stratification, § - incomplete data, na - data not available

**Extended Data Table 2: Diagnostic data of male non-osteoporotic, osteopenic, and osteoporotic individuals.**
Data are given as n, mean ± standard deviation, overall percentages, or percentages of individuals presenting with more than one condition. Differences between study groups were analyzed using a Brown-Forsythe and Welch ANOVA with Dunnett's post-test for numerical data or Fisher's exact test with Bonferroni correction for binary variables (p<0.05 vs. CON-M, CON-H, or CON-F).

| Study groups | CON-H | CON-F | OPE | OPO | CON-M |
|---|---|---|---|---|---|
| Study center | Charité – Universitäts-medizin Berlin | Ruhr-Universität Bochum | | | Charité – Universitäts-medizin Berlin |
| **General** | | | | | |
| n male | 7 | 8 | 8 | 19 | 4 |
| Age [years] | 72.6 ± 7.1 | 65.0 ± 10.1 | 70.1 ± 12.9 | 73.7 ± 11.6 | 33.8 ± 8.9 |
| *vs. CON-M* | *p=0.002* | *p=0.003* | *p=0.001* | *p=0.002* | |
| *vs. CON-H* | | *p=0.370* | *p=0.980* | *p=0.997* | *p=0.002* |
| *vs. CON-F* | *p=0.370* | | *p=0.853* | *p=0.243* | *p=0.003* |
| BMI [kg/m²] | 23.1 ± 3.1 | 26.6 ± 5.4 | 27.3 ± 2.4 | 24.5 ± 3.3 | na |
| *vs. CON-H* | | *p=0.363* | *p=0.041* | *p=0.719* | |
| *vs. CON-F* | *p=0.363* | | *p=0.981* | *p=0.664* | |
| **Osteoporosis and frailty assessment** | | | | | |
| BMD [T score] | no known osteoporosis | -0.2 ± 1.0 | -1.8 ± 0.5 | -3.2 ± 0.5 | no known osteoporosis |
| *vs. CON-F* | | | *p=0.003* | *p<0.001* | |
| ≥ 1 Pre-existing orthopedic condition§ [%] | 0.0 | 100.0 | 62.5 | 31.6 | na |
| *vs. CON-H* | | *p=0.018* | *p=0.545* | *p>0.999* | |
| *vs. CON-F* | *p=0.018* | | *p=0.600* | *p=0.006* | |
| | (n = 3/7) | (n = 8/8) | (n = 8/8) | (n = 19/19) | |
| ≥ 1 Osteochondrosis & Disc prolapse§ [%] | 0.0 (n = 4/7) | 12.5 (n = 8/8) | 25.0 (n = 8/8) | 5.3 (n = 19/19) | na |
| *vs. CON-H* | | *p>0.999* | *p>0.999* | *p>0.999* | |
| *vs. CON-F* | *p>0.999* | | *p>0.999* | *p>0.999* | |
| Rheumatoid arthritis [%] | 0.0 | 25.0 | 12.5 | 5.3 | na |
| *vs. CON-H* | | *p>0.999* | *p>0.999* | *p>0.999* | |
| *vs. CON-F* | *p>0.999* | | *p>0.999* | *p=0.603* | |
| ≥ 1 Fall within last year§ [%] | 16.7 | 50.0 | 25.0 | 68.4 | na |
| *vs. CON-H* | | *p=0.902* | *p>0.999* | *p=0.168* | |
| *vs. CON-F* | *p=0.902* | | *p>0.999* | *p>0.999* | |
| | (n = 6/7) | (n = 8/8) | (n = 8/8) | (n = 19/19) | |
| Difficulty climbing stairs [%] | 0.0 | 62.5 | 50.0 | 63.2 | na |
| *vs. CON-H* | | *p=0.077* | *p=0.231* | *p=0.019* | |
| *vs. CON-F* | *p=0.077* | | *p>0.999* | *p>0.999* | |
| ≥ 1 ∑ numerical frailty assessment [%] | 14.3 | 100.0 | 100.0 | 84.2 | na |
| *vs. CON-H* | | *p=0.004* | *p=0.004* | *p=0.007* | |
| *vs. CON-F* | *p=0.004* | | *p>0.999* | *p>0.999* | |
| HGS/BMI§ | 1.837 ± 0.83 | 1.603 ± 0.80 | 1.445 ± 0.66 | 1.386 ± 0.30 | na |
| *vs. CON-H* | | *p=0.932* | *p=0.721* | *p=0.530* | |
| *vs. CON-F* | *p=0.932* | | *p=0.962* | *p=0.839* | |
| | (n = 6/7) | (n = 8/8) | (n = 8/8) | (n = 18/19) | |

**Continuation Extended Data Table 2**

| Study groups | CON-H | CON-F | OPE | OPO | CON-M |
|---|---|---|---|---|---|
| **Clinical laboratory diagnostics** | | | | | |
| Calcium [mmol/l] | na | 2.21 ± 0.11 | 2.32 ± 0.11 | 2.20 ± 0.14 | na |
| *vs. CON-F* | | | *p=0.156* | *p=0.988* | |
| Phosphate [mg/dl] | na | 2.99 ± 0.60 | 3.21 ± 0.38 | 3.16 ± 0.53 | na |
| *vs. CON-F* | | | *p=0.615* | *p=0.739* | |
| ALP [IU/l] | na | 70.9 ± 29.9 | 84.0 ± 22.4 | 100.3 ± 34.3 | na |
| *vs. CON-F* | | | *p=0.553* | *p=0.080* | |
| VitD§ [ng/ml] | na | 20.8 ± 11.4 | 28.0 ± 13.4 | 17.2 ± 12.1 | na |
| *vs. CON-F* | | | *p=0.447* | *p=0.733* | |
| | | (n = 8/8) | (n = 8/8) | (n = 16/19) | |
| LDL-C§ [mg/dl] | na | 101.0 ± 46.0 | 117.1 ± 34.7 | 96.1 ± 37.0 | na |
| *vs. CON-F* | | | *p=0.681* | *p=0.956* | |
| | | (n = 8/8) | (n = 8/8) | (n = 17/19) | |
| HDL-C§ [mg/dl] | na | 50.0 ± 15.6 | 49.0 ± 10.7 | 51.9 ± 16.2 | na |
| *vs. CON-F* | | | *p=0.986* | *p=0.952* | |
| | | (n = 8/8) | (n = 8/8) | (n = 17/19) | |
| **Cardiovascular assessment** | | | | | |
| IMT§ [mm] | na | 0.850 ± 0.28 | 0.871 ± 0.18 | 0.936 ± 0.14 | na |
| *vs. CON-F* | | | *p=0.913* | *p=0.097* | |
| | | (n = 6/8) | (n = 7/8) | (n = 11/19) | |
| PWV§ [m/s] | na | 7.15 ± 3.75 | 7.70 ± 4.42 | 8.97 ± 2.53 | na |
| *vs. CON-F* | | | *p>0.999* | *p=0.947* | |
| | | (n = 2/8) | (n = 5/8) | (n = 10/19) | |
| pSBP [mm Hg] | na | 128.4 ± 20.9 | 122.0 ± 20.5 | 132.7 ± 20.9 | na |
| *vs. CON-F* | | | *p=0.789* | *p=0.857* | |
| pDBP [mm Hg] | na | 81.3 ± 18.6 | 74.0 ± 10.6 | 75.3 ± 12.0 | na |
| *vs. CON-F* | | | *p=0.579* | *p=0.654* | |
| **Previous cardiovascular diagnoses** | | | | | |
| Hypertension§ [%] | 75.0 | 75.0 | 37.5 | 73.7 | na |
| *vs. CON-H* | | *p>0.999* | *p>0.999* | *p>0.999* | |
| *vs. CON-F* | *p>0.999* | | *p=0.944* | *p>0.999* | |
| | (n = 4/7) | (n = 8/8) | (n = 8/8) | (n = 19/19) | |
| Stroke [%] | 0.0 | 12.5 | 25.0 | 31.6 | na |
| *vs. CON-H* | | *p>0.999* | *p>0.999* | *p=0.437* | |
| *vs. CON-F* | *p>0.999* | | *p>0.999* | *p>0.999* | |
| Coronary artery disease [%] | 0.0 | 37.5 | 25.0 | 26.3 | na |
| *vs. CON-H* | | *p=0.600* | *p>0.999* | *p=0.835* | |
| *vs. CON-F* | *p=0.600* | | *p>0.999* | *p>0.999* | |
| Cardiac infarction [%] | 0.0 | 12.5 | 0.0 | 5.3 | na |
| *vs. CON-H* | | *p>0.999* | *p>0.999* | *p>0.999* | |
| *vs. CON-F* | *p>0.999* | | *p>0.999* | *p>0.999* | |
| Cardiac insufficiency [%] | 0.0 | 12.5 | 25.0 | 5.3 | na |
| *vs. CON-H* | | *p>0.999* | *p>0.999* | *p>0.999* | |
| *vs. CON-F* | *p>0.999* | | *p>0.999* | *p>0.999* | |

**Continuation Extended Data Table 2**

| Study groups | CON-H | CON-F | OPE | OPO | CON-M |
|---|---|---|---|---|---|
| **Continuation previous cardiovascular diagnoses** | | | | | |
| Peripheral artery disease [%] | 0.0 | 12.5 | 0.0 | 0.0 | na |
| *vs. CON-H* | | *p>0.999* | *p>0.999* | *p>0.999* | |
| *vs. CON-F* | *p>0.999* | | *p>0.999* | *p=0.889* | |
| ≥ 1 ∑ previous cardio-vascular diagnoses [%] | 42.9 | 87.5 | 50.0 | 78.9 | na |
| *vs. CON-H* | | *p=0.357* | *p>0.999* | *p=0.447* | |
| *vs. CON-F* | *p=0.357* | | *p=0.846* | *p>0.999* | |
| **Assessment of secondary diagnoses and related medication** | | | | | |
| Diabetes [%] | 0.0 | 0.0 | 0.0 | 15.8 | na |
| *vs. CON-H* | | *p>0.999* | *p>0.999* | *p>0.999* | |
| *vs. CON-F* | *p>0.999* | | *p>0.999* | *p>0.999* | |
| Use of blood sugar reducing medication [%] | 0.0 | 0.0 | 0.0 | 10.5 | na |
| *vs. CON-H* | | *p>0.999* | *p>0.999* | *p>0.999* | |
| *vs. CON-F* | *p>0.999* | | *p>0.999* | *p>0.999* | |
| Use of statins [%] | 14.3 | 37.5 | 0.0 | 42.1 | na |
| *vs. CON-H* | | *p>0.999* | *p>0.999* | *p>0.999* | |
| *vs. CON-F* | *p>0.999* | | *p=0.600* | *p>0.999* | |
| Hyperlipidemia [%] | na | 37.5 | 12.5 | 42.1 | na |
| *vs. CON-F* | | | *p>0.999* | *p>0.999* | |
| Hyperuricemia [%] | na | 12.5 | 0.0 | 5.3 | na |
| *vs. CON-F* | | | *p>0.999* | *p>0.999* | |
| Renal insufficiency [%] | na | 0.0 | 0.0 | 10.5 | na |
| *vs. CON-F* | | | *p>0.999* | *p>0.999* | |
| Current/previous smoker§ [%] | 50.0 | 25.0 | 25.0 | 31.6 | na |
| *vs. CON-H* | | *p>0.999* | *p>0.999* | *p>0.999* | |
| *vs. CON-F* | *p>0.999* | | *p>0.999* | *p>0.999* | |
| | (n = 4/7) | (n = 8/8) | (n = 8/8) | (n = 19/19) | |

CON-H - non-osteoporotic older healthy controls, CON-F - non-osteoporotic frail controls, OPE - osteopenic frail individuals, OPO - osteoporotic frail individuals, CON-M - non-osteoporotic middle-aged controls, n - number, BMD - bone mineral density, BMI - body mass index, ALP - alkaline phosphatase, VitD - (25)-Hydroxy-Vitamin D, LDL-C - low-density lipoprotein cholesterol, HDL-C - high-density lipoprotein cholesterol, no known osteoporosis - self-declaration corresponding to past medical examinations, pSBP - peripheral systolic blood pressure, pDBP - peripheral diastolic blood pressure, IMT - intima-media thickness, PWV - pulse wave velocity, excluded - according to the exclusion criterion applied during stratification, § - incomplete data, na - data not available

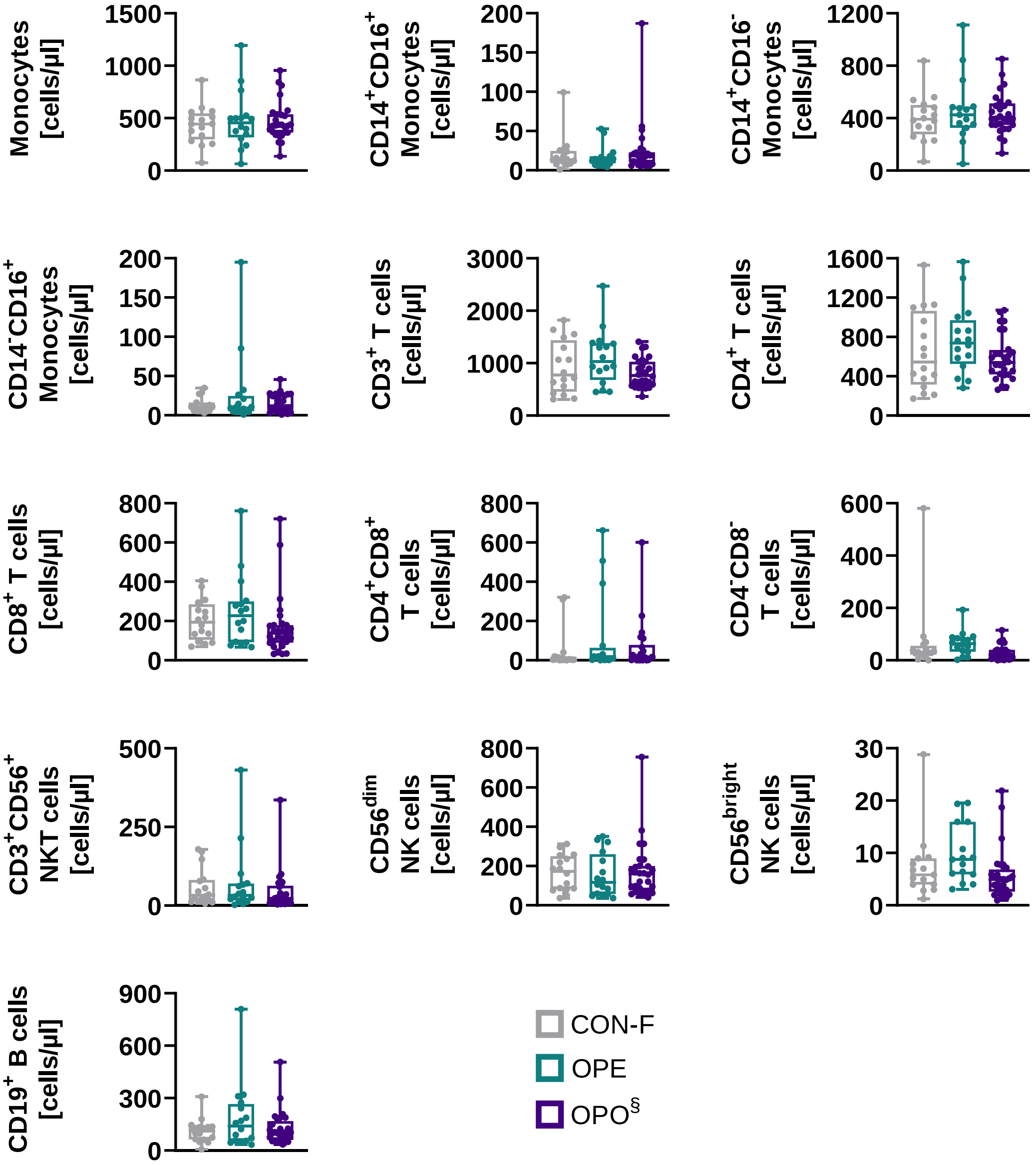


**Extended Data Fig. 1: Monocytes, T cells, natural killer, and B cells do not vary with bone mineral density.**

Flow cytometry analysis of peripheral immune cell subsets in non-osteoporotic frail controls (CON-F; n=16), osteopenic frail individuals (OPE; n=16), and osteoporotic frail individuals (OPO; n=28 out of 38) from the Ruhr-Universität Bochum study center. Differences in immune cell subsets were assessed using Brown-Forsythe and Welch ANOVA with Dunnett's T3 multiple comparisons test vs. CON-F if data were normally distributed. Non-normal data were analyzed with a Kruskal-Wallis test and Dunn's multiple comparison. § indicates incomplete immune cell subset data. No significant differences ($p<0.05$) or distinct trends with $p<0.1$ were observed.

**Extended Data Table 3: Granulocyte-to-lymphocyte ratios are higher in osteoporotic individuals.** Peripheral immune cell subsets of non-osteoporotic frail controls (CON-F), osteopenic frail individuals (OPE), as well as osteoporotic frail individuals (OPO) from the Ruhr-Universität Bochum study center were determined by flow cytometry and normalized to the total lymphocyte count of each individual. The systemic inflammation index (SII) was calculated as the product of the neutrophil-to-lymphocyte ratio (NLR) and platelet count from clinical laboratory measurements. Ratios are given as group means. Differences in immune cell ratios were assessed using Brown-Forsythe and Welch ANOVA with Dunnett's T3 multiple comparisons vs. CON-F ($p<0.05$) for normally distributed data. Non-normal data were analyzed with a Kruskal-Wallis test and Dunn's multiple comparison, $^{\S}$ - indicates incomplete data.

| | **CON-F** | **OPE** | **OPO$^{\S}$** | **p-value** | |
|---|---|---|---|---|---|
| | (n=16/16) | (n=16/16) | (n=28/38) | CON-F vs. OPE | CON-F vs. OPO |
| Granulocytes/Lymphocytes | 3.129 | 2.983 | 4.794 | 0.871 | 0.129 |
| Basophils/Lymphocytes | 0.039 | 0.040 | 0.053 | 0.549 | 0.070 |
| Eosinophils/Lymphocytes | 0.109 | 0.137 | 0.195 | 0.693 | 0.010 |
| Neutrophils/Lymphocytes | 2.981 | 2.807 | 4.546 | 0.703 | 0.218 |
| Monocytes/Lymphocytes | 0.395 | 0.376 | 0.423 | 0.505 | 0.518 |
| $CD14^{+}CD16^{+}$ Monocytes/Lymphocytes | 0.021 | 0.015 | 0.020 | 0.383 | 0.885 |
| $CD14^{+}CD16^{-}$ Monocytes/Lymphocytes | 0.364 | 0.339 | 0.389 | 0.531 | 0.600 |
| $CD14^{-}CD16^{+}$ Monocytes/Lymphocytes | 0.010 | 0.023 | 0.014 | >0.999 | 0.403 |
| $CD3^{+}$ Lymphocytes/Lymphocytes | 0.700 | 0.715 | 0.693 | 0.864 | 0.956 |
| $CD4^{+}$ T cells/Lymphocytes | 0.481 | 0.507 | 0.512 | 0.650 | 0.558 |
| $CD8^{+}$ T cells/Lymphocytes | 0.166 | 0.150 | 0.136 | >0.999 | 0.159 |
| $CD4^{+}CD8^{+}$ T cells/Lymphocytes | 0.069 | 0.075 | 0.053 | 0.813 | 0.952 |
| $CD4^{-}CD8^{-}$ T cells/Lymphocytes | 0.071 | 0.049 | 0.025 | 0.594 | 0.291 |
| $CD3^{+}CD56^{+}$ NKT cells/Lymphocytes | 0.039 | 0.038 | 0.036 | >0.999 | 0.367 |
| $CD56^{dim}$ NK cells/Lymphocytes | 0.141 | 0.106 | 0.135 | 0.369 | 0.967 |
| $CD56^{bright}$ NK cells/Lymphocytes | 0.005 | 0.007 | 0.005 | 0.362 | 0.993 |
| $CD19^{+}$ B cells/Lymphocytes | 0.091 | 0.110 | 0.108 | 0.543 | 0.504 |
| Systemic inflammation index (SII) | 722 | 772 | 1495 | >0.999 | 0.016 |

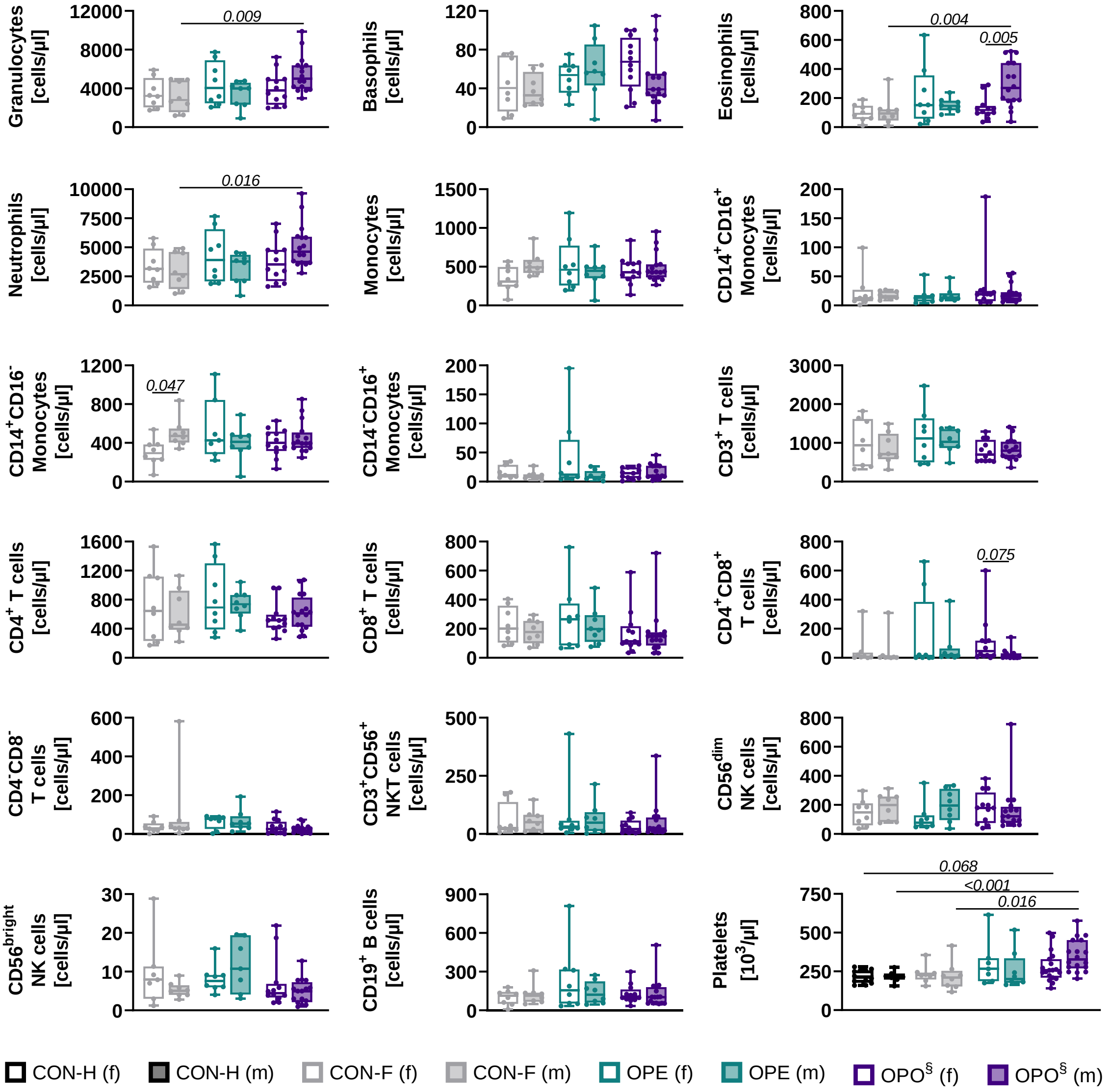


**Extended Data Fig. 2: Eosinophil and neutrophil counts are higher in male osteoporotic individuals, while platelet counts are elevated in both sexes.**

Flow cytometry analysis of peripheral immune cell subsets in female (f) and male (m) non-osteoporotic frail controls (CON-F; n=8/group), osteopenic frail individuals (OPE; n=8/group), and osteoporotic frail individuals (OPO; f: n=12 out of 19; m: n=16 out of 19), as well as platelet counts in non-osteoporotic healthy older controls (CON-H; f: n=12; m: n=7). Differences in immune cell subsets were assessed using Brown-Forsythe and Welch ANOVA with Dunnett's T3 multiple comparisons vs. CON-F or CON-H for normally distributed data. Non-normally distributed data were analyzed with Kruskal-Wallis and Dunn's multiple comparison tests. Only significant differences (p<0.05) and distinct trends with p<0.1 are shown; § - indicates incomplete data.

**Extended Data Table 4: Eosinophil-to-lymphocyte ratios are higher in male osteoporotic individuals, while neutrophil-to-lymphocyte ratios are elevated in both sexes.**
Peripheral granulocytes of female and male non-osteoporotic frail controls (CON-F), osteopenic frail individuals (OPE), and osteoporotic frail individuals (OPO) from the Ruhr-Universität Bochum study center were determined by flow cytometry and normalized to the total lymphocyte count of each individual. The systemic inflammation index (SII) was calculated as the product of the neutrophil-to-lymphocyte ratio (NLR) and platelet count from clinical laboratory measurements. Ratios are given as group means per sex. Differences among study groups of the same sex and between females and males within one group were assessed using Brown-Forsythe and Welch ANOVA with Dunnett's T3 multiple comparisons test ($p<0.05$), § - indicates incomplete data.

| | **CON-F** | **OPE** | **OPO§** | **p-values (among study groups)** | |
|---|---|---|---|---|---|
| | | **female** | | | |
| | (n=8/8) | (n=8/8) | (n=12/19) | CON-F vs. OPE | CON-F vs. OPO |
| Granulocytes/Lymphocytes | 3.017 | 3.701 | 4.116 | 0.830 | 0.375 |
| Basophils/Lymphocytes | 0.041 | 0.041 | 0.060 | 0.998 | 0.494 |
| Eosinophils/Lymphocytes | 0.106 | 0.167 | 0.112 | 0.683 | 0.990 |
| Neutrophils/Lymphocytes | 2.870 | 3.493 | 3.994 | 0.856 | 0.380 |
| Systemic inflammation index | 646 | 997 | 1159 | 0.438 | 0.190 |
| | | **male** | | | |
| | (n=8/8) | (n=8/8) | (n=15/19) | CON-F vs. OPE | CON-F vs. OPO |
| Granulocytes/Lymphocytes | 3.240 | 2.266 | 5.303 | 0.415 | 0.127 |
| Basophils/Lymphocytes | 0.037 | 0.039 | 0.048 | 0.976 | 0.704 |
| Eosinophils/Lymphocytes | 0.112 | 0.107 | 0.258 | 0.991 | 0.037 |
| Neutrophils/Lymphocytes | 3.091 | 2.120 | 4.997 | 0.400 | 0.153 |
| Systemic inflammation index | 798 | 575 | 1763 | 0.699 | 0.043 |
| | | **p-values (female vs. male of the study group)** | | | |
| Granulocytes/Lymphocytes | 0.991 | 0.571 | 0.602 | | |
| Basophils/Lymphocytes | 0.997 | 0.997 | 0.713 | | |
| Eosinophils/Lymphocytes | >0.999 | 0.738 | 0.004 | | |
| Neutrophils/Lymphocytes | 0.990 | 0.606 | 0.681 | | |
| Systemic inflammation index | 0.916 | 0.478 | 0.379 | | |

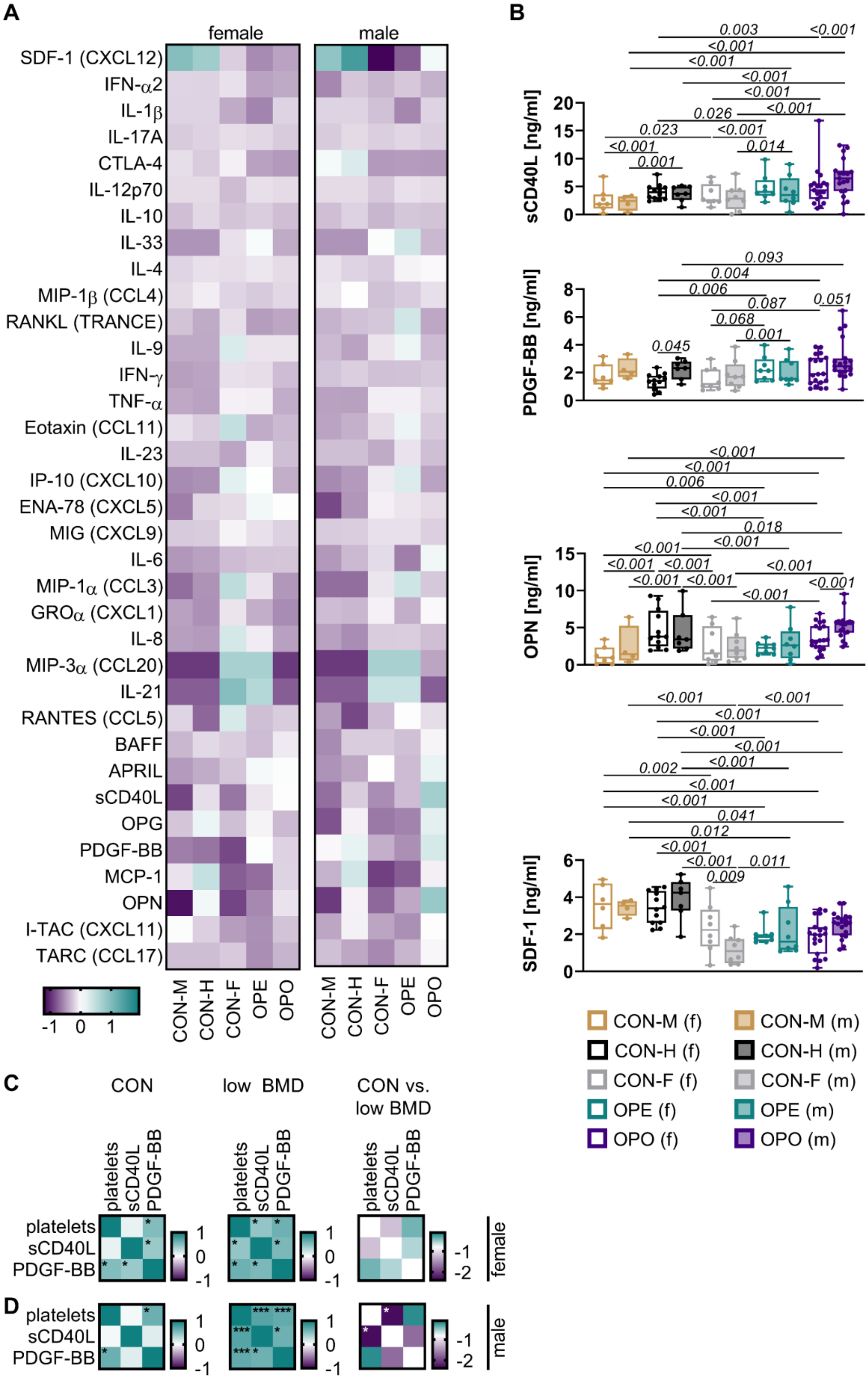
A
female
male
SDF-1 (CXCL12)
IFN-α2
IL-1β
IL-17A
CTLA-4
IL-12p70
IL-10
IL-33
IL-4
MIP-1β (CCL4)
RANKL (TRANCE)
IL-9
IFN-γ
TNF-α
Eotaxin (CCL11)
IL-23
IP-10 (CXCL10)
ENA-78 (CXCL5)
MIG (CXCL9)
IL-6
MIP-1α (CCL3)
GROα (CXCL1)
IL-8
MIP-3α (CCL20)
IL-21
RANTES (CCL5)
BAFF
APRIL
sCD40L
OPG
PDGF-BB
MCP-1
OPN
I-TAC (CXCL11)
TARC (CCL17)
-1 0 1
CON-M
CON-H
CON-F
OPE
OPO
B
sCD40L [ng/ml]
0.003
<0.001
<0.001
<0.001
<0.001
<0.001
<0.001
0.026
0.023
<0.001
<0.001
0.014
0.001
PDGF-BB [ng/ml]
0.093
0.004
0.006
0.087
0.051
0.068
0.001
0.045
OPN [ng/ml]
<0.001
<0.001
0.006
<0.001
<0.001
0.018
<0.001
<0.001
<0.001
<0.001
<0.001
<0.001
<0.001
<0.001
<0.001
<0.001
SDF-1 [ng/ml]
<0.001
<0.001
<0.001
<0.001
<0.001
0.002
<0.001
<0.001
<0.001
0.041
0.012
<0.001
<0.001
0.011
0.009
CON-M (f)
CON-M (m)
CON-H (f)
CON-H (m)
CON-F (f)
CON-F (m)
OPE (f)
OPE (m)
OPO (f)
OPO (m)
C
CON
low BMD
CON vs. low BMD
platelets
sCD40L
PDGF-BB
female
D
male

**Extended Data Fig. 3: Sex differences in osteoporotic and osteopenic cytokine and chemokine patterns.**

Screening of sera from non-osteoporotic middle-aged controls (CON-M), non-osteoporotic healthy older controls (CON-H), non-osteoporotic frail controls (CON-F), osteopenic (OPE), and osteoporotic (OPO) individuals using bead-based multiplex assays. **(A)** Clustered expression levels of female (f) and male (m) individuals, shown as median Z-scores. **(B)** Factors with significantly different expression levels identified by overall group analysis (female and male, Figure 2). Differences among study groups of the same sex were assessed using two-way ANOVAs with Dunnett's post-test vs. controls (CON-M, CON-H, CON-F). Differences between females and males within one study group were assessed using a Sidak post-test. Only significant differences ($p<0.05$) and distinct trends ($p<0.1$) are shown. Spearman correlation of platelet counts, sCD40L, and PDGF-BB levels in **(C)** female and **(D)** male non-osteoporotic controls (CON: CON-M, CON-H, CON-F) and in osteopenic and osteoporotic individuals (low BMD: OPE, OPO). Differences between correlation coefficients of CON and low BMD within the same sex were assessed using Fisher's Z-tests; * $p<0.05$, ** $p<0.001$, *** $p<0.0001$.

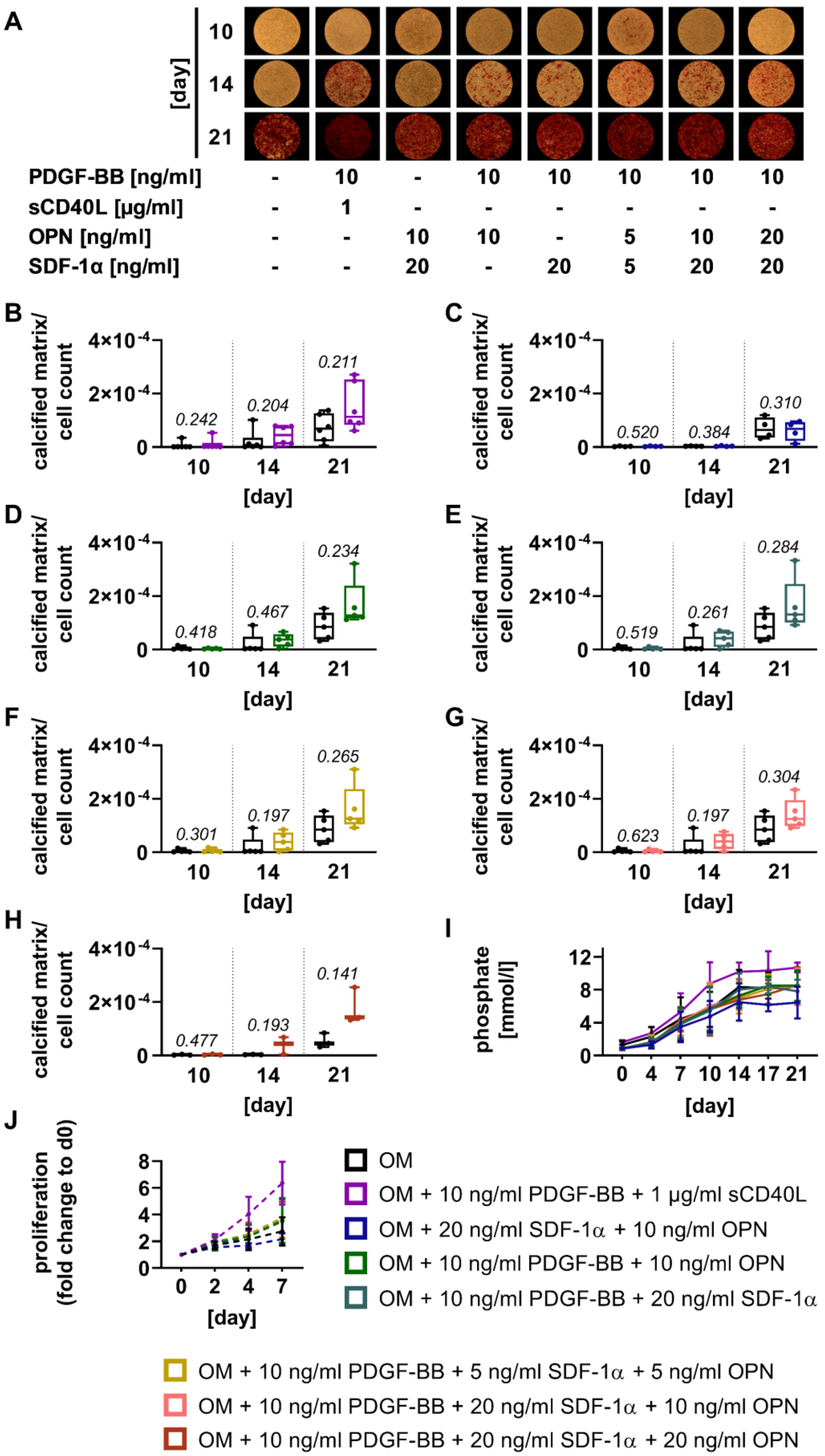
A
[day]
10
14
21
PDGF-BB [ng/ml] - 10 - 10 10 10 10 10
sCD40L [µg/ml] - 1 - - - - - -
OPN [ng/ml] - - 10 10 - 5 10 20
SDF-1α [ng/ml] - - 20 - 20 5 20 20
B
C
D
E
F
G
H
calcified matrix/ cell count
4×10-4
2×10-4
0
[day]
0.242
0.204
0.211
0.520
0.384
0.310
0.418
0.467
0.234
0.519
0.261
0.284
0.301
0.197
0.265
0.623
0.197
0.304
0.477
0.193
0.141
I
phosphate [mmol/l]
0 4 7 10 14 17 21
J
proliferation (fold change to d0)
0 2 4 7
OM
OM + 10 ng/ml PDGF-BB + 1 µg/ml sCD40L
OM + 20 ng/ml SDF-1α + 10 ng/ml OPN
OM + 10 ng/ml PDGF-BB + 10 ng/ml OPN
OM + 10 ng/ml PDGF-BB + 20 ng/ml SDF-1α
OM + 10 ng/ml PDGF-BB + 5 ng/ml SDF-1α + 5 ng/ml OPN
OM + 10 ng/ml PDGF-BB + 20 ng/ml SDF-1α + 10 ng/ml OPN
OM + 10 ng/ml PDGF-BB + 20 ng/ml SDF-1α + 20 ng/ml OPN

**Extended Data Fig. 4: Combinations with PDGF-BB increase SMC calcification and proliferation.** SMC undergoing osteogenic differentiation (OM - osteogenic medium) in an *in vitro* model of SMC calcification were stimulated with combinations of recombinant PDGF-BB, sCD40L, OPN, and SDF-1α. Deposited calcified matrix was stained with alizarin red. **(A**) Representative images of the deposited calcified matrix and **(B-H)** quantified staining normalized to cell count. A paired, two-tailed t-test vs. OM without supplements was used to assess significant differences in SMC calcification at each time point ($p<0.05$). **(I)** Phosphate levels determined in the supernatant during osteogenic differentiation. For statistical analysis of phosphate levels at each time point, a mixed-effects model with Geisser-Greenhouse correction and Dunnett's post-test vs. OM was used ($p<0.05$). No significant differences were detected. **(J)** Proliferation of SMC after stimulation with the respective combinations of recombinant factors in expansion medium (EM; dashed line). For statistical analysis at each time point, a two-way ANOVA with Geisser-Greenhouse correction and Dunnett's post-test vs. EM without supplements was used ($p<0.05$). No significant differences were detected.

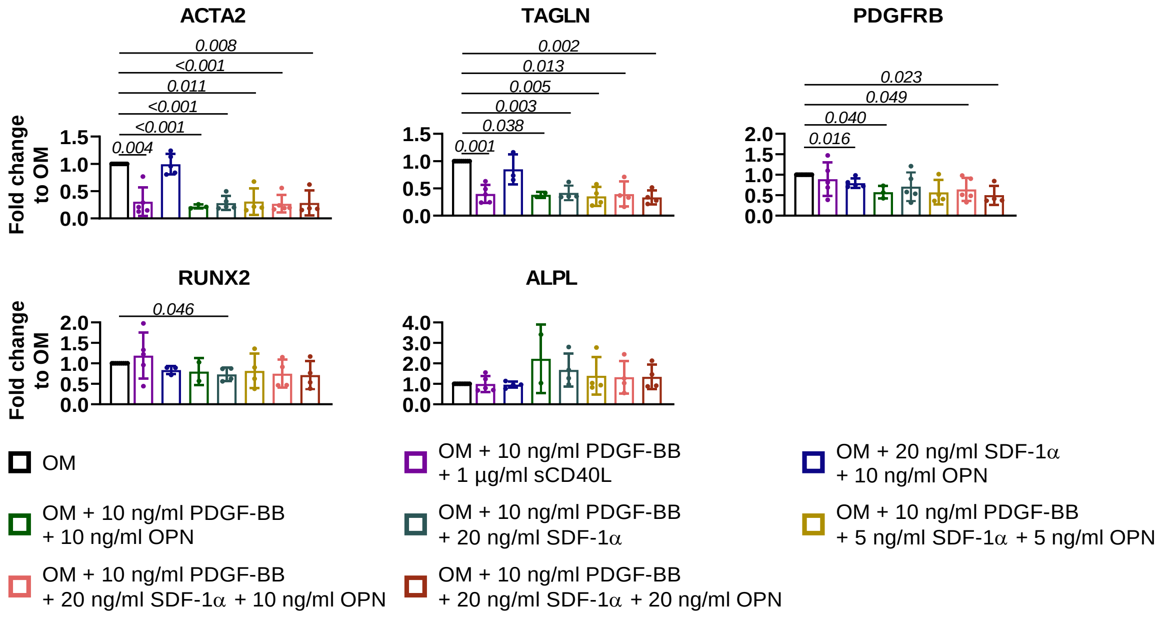


**Extended Data Fig. 5: Combinations with PDGF-BB decrease SMC marker expression.** Gene expression analysis of SMC after four days of osteogenic differentiation (OM - osteogenic medium) with the respective combinations of recombinant factors. A paired, two-tailed t-test vs. OM was used to assess significant differences in independent experiments ($p<0.05$). Only significant differences are shown.

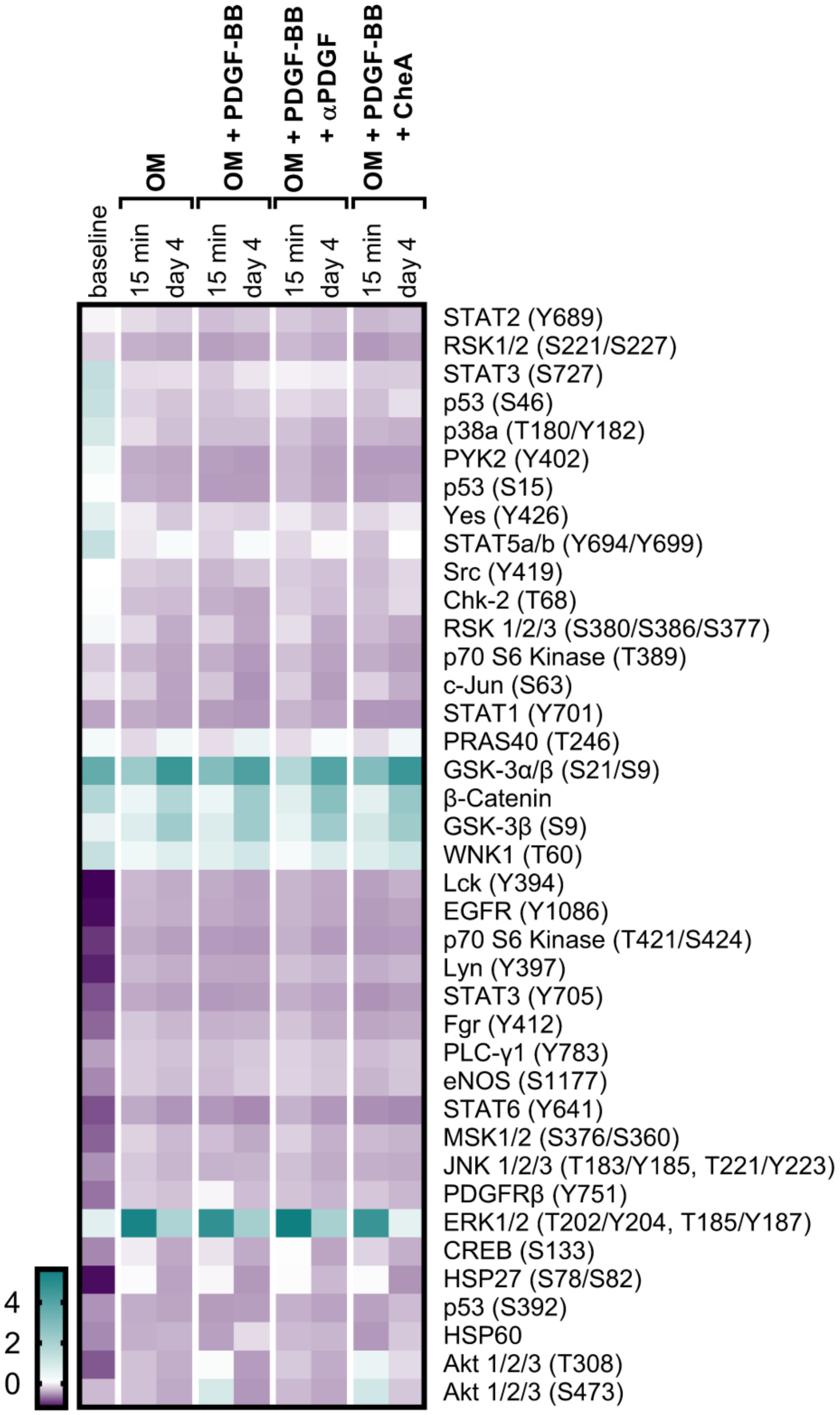


**Extended Data Fig. 6: Chebulinic acid reduces pro-osteogenic p-ERK signaling in SMC to baseline levels after four days of osteogenic differentiation with PDGF-BB.**
Kinase phosphorylation profiles in human coronary artery SMC from one healthy donor after 24 h starvation (baseline) and stimulation with osteogenic medium (OM) supplemented with 10 ng/ml PDGF-BB and either 25 µg/ml neutralizing polyclonal PDGF antibody (αPDGF) or 50 µM chebulinic acid (CheA) for 15 min and four days. Clustered phosphorylation levels are given as Z-scores.
