## Supplemental Material for "A complex osteoporotic milieu is associated with arterial stiffening and PDGF-BB-mediated calcification of human smooth muscle cells"

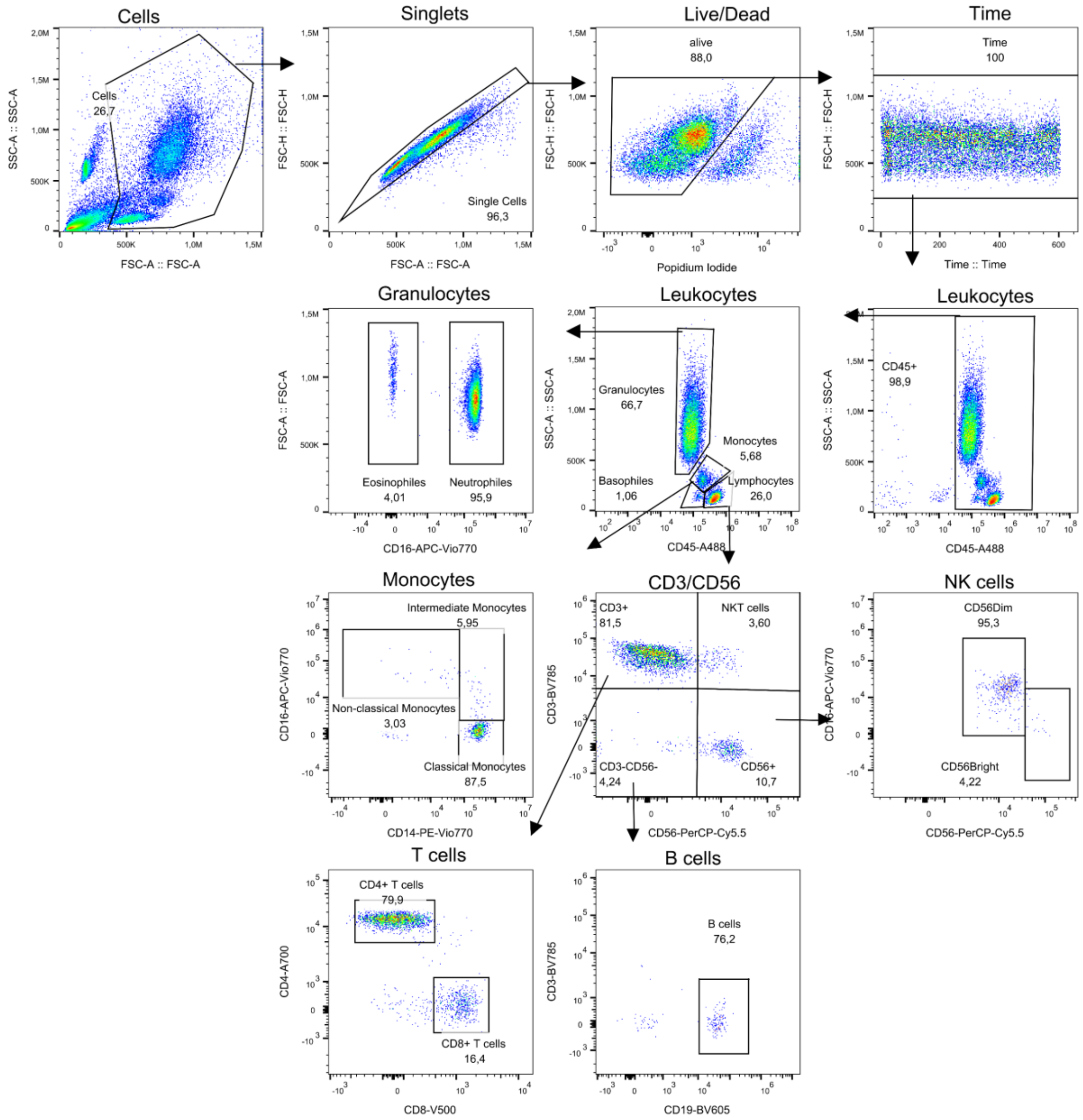


**Supplementary Fig. 1: Gating strategy for identification of peripheral immune cell subsets.** Peripheral immune cell subsets of individuals recruited at the Ruhr-Universität Bochum study center were analyzed by flow cytometry using whole blood. Cells were identified based on forward and side scatter profiles, doublets were excluded, and living cells were gated as propidium iodide-negative. Granulocytes, monocytes, and lymphocytes were distinguished by CD45 expression and side scatter profile. Granulocytes were separated by CD16 into eosinophils ($CD16^-$) and neutrophils ($CD16^+$). Monocytes were subdivided according to CD14 and CD16 expression into classical ($CD14^{++}CD16^-$), intermediate ($CD14^{++}CD16^+$), and non-classical ($CD14^+CD16^+$) monocytes. Lymphocytes were separated by CD3 and CD56 into NK ($CD3^-CD56^+$), NKT ($CD3^+CD56^+$), and T cells ($CD3^+CD56^-$). T cells were further distinguished into $CD4^+$ helper and $CD8^+$ cytotoxic T cells. B cells were identified as $CD3^-CD56^-CD19^+$. Fractions of all immune cell populations are given as percentages.

**Supplementary Table 1: Primary, monoclonal antibodies used for flow cytometry analysis.**

| Antigen | Fluorophore | Antibody Type | Clone | Vendor | Titer |
|---|---|---|---|---|---|
| CD3 | BV785 | Mouse IgG2a (Cat# 317329, RRID:AB_11219196) | OKT3 | BioLegend® | 1:200 |
| CD4 | A700 | Mouse IgG2b (Cat# 317425, RRID:AB_571942) | OKT4 | BioLegend® | 1:200 |
| CD8 | V500 | Mouse IgG1 (Cat# 560774, RRID:AB_1937325) | RPA-T8 | BD | 1:25 |
| CD14 | PE-Vio770 | Mouse IgG2a (Cat# 170-078-012) | TÜK4 | Miltenyi Biotec | 1:100 |
| CD16 | APC-Vio770 | recombinant human IgG1 (Cat# 170-078-088) | REA423 | Miltenyi Biotec | 1:100 |
| CD19 | BV605 | Mouse IgG1 (Cat# 302243, RRID:AB_2562014) | HIB19 | BioLegend® | 1:200 |
| CD45 | A488 | Mouse IgG1 (Cat# 368535, RRID:AB_2721363) | 2D1 | BioLegend® | 1:200 |
| CD56 | PerCP-Cy5.5 | Mouse IgG1 (Cat# 318321, RRID:AB_893391) | HCD56 | BioLegend® | 1:50 |
| HLA-DR | BV650 | Mouse IgG2a (Cat# 307649, RRID:AB_2562544) | L243 | BioLegend® | 1:100 |
| Propidium Iodide | - | - | - | Thermo Fisher | 1 µg/ml |

**Supplementary Table 2: List of primers used for qRT-PCR.**

| Gene | Abbreviation | Direction | Sequences 5′ → 3′ |
|---|---|---|---|
| Alkaline phosphatase, tissue-nonspecific | ALPL | Forward | ATGTTCCTGGGAGATGGGATG |
| | | Reverse | ACCTGGGCATTGGTGTTGTA |
| Aortic smooth muscle actin, alpha2 | ACTA2 | Forward | GCCAAGCACTGTCAGGAATC |
| | | Reverse | GGGTACTTCAGGGTCAGGAT |
| Platelet-derived growth factor receptor β | PDGFRB | Forward | TCTTTGTGCCAGATCCCACC |
| | | Reverse | AGTGCAACGTCCCCTTTCTT |
| Ribosomal protein L13a | RPL13A | Forward | CCTGGAGGAGAAGAGGAAAGAGA |
| | | Reverse | TTGAGGACCTCTGTGTATTTGTCAA |
| Runt-related transcription factor 2 | RUNX2 | Forward | CTCCTACCTGAGCCAGATGA |
| | | Reverse | CGGGGTGTAAGTAAAGGTGG |
| Transgelin | TAGLN | Forward | GAAACCCACCCTCTCAGTCA |
| | | Reverse | ATGTCTGGGGAAAGCTCCT |
