## Supplementary material for "A complex osteoporotic milieu is associated with arterial stiffening and PDGF-BB-mediated calcification of human smooth muscle cells": STROBE Checklist

STROBE Statement—checklist of items that should be included in reports of observational studies

| | Item No. | Recommendation | Page No. | Relevant text from manuscript |
|---|---|---|---|---|
| **Title and abstract** | 1 | (*a*) Indicate the study's design with a commonly used term in the title or the abstract | Abstract 2 | In the abstract the study is indicated as "retrospective exploratory study" |
| | | (*b*) Provide in the abstract an informative and balanced summary of what was done and what was found | Abstract 2 | The abstract provides a comprehensive summary of the study approach and its main findings. Please see the corresponding text in the main manuscript. |
| **Introduction** | | | | |
| Background/ rationale | 2 | Explain the scientific background and rationale for the investigation being reported | Introduction 3, 4 | The Introduction outlines the observed link between osteoporosis and vascular calcification:<br>"Increasing clinical and epidemiological evidence demonstrates a distinct association between osteoporosis and vascular calcification."<br><br>gives an overview of factors investigated in the past regarding a bone-vascular interplay:<br>"Recent murine studies have identified platelet-derived growth factor beta (PDGF-BB), secreted by preosteoclasts in the bone marrow, as a key mediator of arterial stiffening and vascular calcification in the brain. Earlier studies, moreover, established matrix Gla protein as an important paracrine inhibitor of calcification in both bone and vasculature in mice and human atherosclerotic lesions. Furthermore, osteopontin (OPN) is secreted by activated osteoclasts and promotes mineralization in human bone, yet it has also been reported to inhibit arterial calcification.",<br>"As osteoporosis, atherosclerosis, and vascular calcification are driven by inflammatory processes, an inflammatory link between both conditions has also been suggested. […]",<br><br>points out that a comprehensive understanding is still lacking:<br>"Independent of shared risk factors such as advanced age, sedentary lifestyle, smoking, menopause, and diabetes, the pathological pathways linking bone and vasculature remain under investigation.",<br><br>and suggests to conceptualize this potential multifactorial interplay as a complex adaptive system to broaden the understanding of the pathological bone-vascular cross-talk:<br>"[…] we conceptualize the pathological bone-vascular cross-talk as a CAS to improve understanding of disease complexity […]." |
| Objectives | 3 | State specific objectives, including any prespecified hypotheses | Introduction 4 | "[…] improve understanding of disease complexity and help to identify novel opportunities for targeted interventions that support healthier aging." |

| | | | | |
|---|---|---|---|---|
| **Methods** | | | | |
| Study design | 4 | Present key elements of study design early in the paper | Results 5, 6<br><br>Methods 17, 18 | The first section of the Results and Fig. 1A describe the outline of the study groups in detail. An overview of the consecutive study design is given in Fig. 1B.<br><br>Additional information on the inclusion and exclusion criteria is provided in the first Methods section "Sample acquisition and study criteria". |
| Setting | 5 | Describe the setting, locations, and relevant dates, including periods of recruitment, exposure, follow-up, and data collection | Methods 17 - 20 | Sample acquisition and study criteria as well as the diagnostic assessment are described in detail in the Methods sections "Sample acquisition and study criteria" and "Diagnostic assessment of osteoporosis, frailty, and risk factors for cardiovascular disease". |
| Participants | 6 | (*a*) *Cohort study*—Give the eligibility criteria, and the sources and methods of selection of participants. Describe methods of follow-up<br>*Case-control study*—Give the eligibility criteria, and the sources and methods of case ascertainment and control selection. Give the rationale for the choice of cases and controls<br>*Cross-sectional study*—Give the eligibility criteria, and the sources and methods of selection of participants | Results 5, 6<br><br>Methods 17, 18 | Stratification criteria of the participants selected for the study are outlined in the first Results section "Study groups and their characteristics indicate links between osteoporosis, frailty, and arterial stiffening" and Fig. 1A.<br>Additional information on the inclusion and exclusion criteria of the studies used for participant selection for the present investigation is provided in the first Methods section "Sample acquisition and study criteria". No follow-up data are available from the recruiting studies used for this investigation. |
| | | (*b*) *Cohort study*—For matched studies, give matching criteria and number of exposed and unexposed<br>*Case-control study*—For matched studies, give matching criteria and the number of controls per case | | Not applicable |
| Variables | 7 | Clearly define all outcomes, exposures, predictors, potential confounders, and effect modifiers. Give diagnostic criteria, if applicable | Results 5, 6<br>Methods 17, 18 | Corresponding information can be found in the Results section "Study groups and their characteristics indicate links between osteoporosis, frailty, and arterial stiffening" and in the Methods section "Sample acquisition and study criteria". |
| Data sources/ measurement | 8* | For each variable of interest, give sources of data and details of methods of assessment (measurement). Describe comparability of assessment methods if there is more than one group | Methods 18 - 20 | Methods of diagnostic assessment as well as comparability of data amongst the study groups are described in the Methods section "Diagnostic assessment of osteoporosis, frailty, and risk factors for cardiovascular disease". |
| Bias | 9 | Describe any efforts to address potential sources of bias | Results 5, 6 | To address the potential biases of age and frailty and to consider the respective heterogeneity of the older population, we included three non-osteoporotic control groups in our retrospective exploratory study: middle-aged non-osteoporotic controls (CON-M), non-osteoporotic healthy older controls (CON-H), non-osteoporotic frail controls (CON-F). Moreover, comparable numbers of female and male participants were selected for the study groups wherever possible. Potential differences between female and male participants were addressed by data analyses according to the participants sex. Additional bias may result from the inclusion of participants with the most complete data and serum samples, which was essential for the breadth of analyses performed. |

| | | | | |
|---|---|---|---|---|
| Study size | 10 | Explain how the study size was arrived at | Results 5, 6 | For this retrospective exploratory study participants were selected according to stratification criteria, potential sources of bias, and availability of diagnostic data and samples. Study groups of comparable size and wherever possible a comparable number of female and male participants were formed. |
| Quantitative variables | 11 | Explain how quantitative variables were handled in the analyses. If applicable, describe which groupings were chosen and why | Methods 28 - 30 | "Differences in group sizes were addressed by reporting percentages or means, and sex-specific analyses were performed where possible. Categorical diagnoses and discrete counts, e.g. comorbidities and number of falls were transformed into binary variables, $\geq$1 diagnosis or fall was coded as "yes" and none as "no", to enable comparability across groups." Differences between the study groups were analyzed using the statistical tests outlined in Methods section "Statistical analysis and correlation" and the Figure and Table legends. Potential differences between female and male participants were addressed by data analysis according to the participants sex. |
| Statistical methods | 12 | (*a*) Describe all statistical methods, including those used to control for confounding | Methods 28 - 30 | Statistical methods are described in the Methods section "Statistical analysis and correlation". |
| | | (*b*) Describe any methods used to examine subgroups and interactions | Methods 28 - 30 | Methods used to examine subgroups and interactions are described in the Methods section "Statistical analysis and correlation". |
| | | (*c*) Explain how missing data were addressed | Figures and Tables | § indicates incomplete data, na - data not available |
| | | (*d*) *Cohort study*—If applicable, explain how loss to follow-up was addressed<br>*Case-control study*—If applicable, explain how matching of cases and controls was addressed<br>*Cross-sectional study*—If applicable, describe analytical methods taking account of sampling strategy | Methods 17<br><br>Results 5, 6 | "The OsteoSys study and the study "Einfluss des Serums auf das regenerative Potential von mesenchymalen Stammzellen (MSC) in vitro" collected data only at the time of enrollment, with no follow-up data available. Accordingly, from the Frailty study, only data and samples obtained at study entry were used."<br>Only data collected in a comparable manner at both recruitment centers were used for comparative analysis in our exploratory study: "Frailty assessments were conducted in all older participants. However, since the recorded items differed between study centers, comparative analysis included only data collected in a comparable manner (number of falls within the last year, difficulty climbing stairs, and hand grip strength (HGS)), […]." |
| | | (*e*) Describe any sensitivity analyses | | Not applicable |
| **Results** | | | | |
| Participants | 13* | (a) Report numbers of individuals at each stage of study—e.g. numbers potentially eligible, examined for eligibility, confirmed eligible, included in the study, completing follow-up, and analyzed | Figures and Tables | The number of individuals included in the analyses are indicated by the respective data points in the Figures of the *ex vivo* analysis. Missing or incomplete data in Figures and Tables are indicated by § symbol. Wherever possible Tables state the number of individuals included in the analysis. |
| | | (b) Give reasons for non-participation at each stage | Figures and Tables | All data available were included in the respective analysis. |
| | | (c) Consider use of a flow diagram | | Not applicable |
| Descriptive data | 14* | (a) Give characteristics of study participants (e.g. demographic, clinical, social) and information on exposures and potential confounders | Results 5, 6 | A detailed characterization of the study participants is given in Results section "Study groups and their characteristics indicate links between osteoporosis, frailty, and arterial stiffening", Table 1 and Extended Data Tables 1 and 2. |
| | | (b) Indicate number of participants with missing data for each variable of interest | Figures and Tables | The number of individuals included in the analysis is indicated by the respective data points in the Figures of the *ex vivo* analysis. Missing or incomplete data in Figures and Tables are indicated by § symbol. Wherever possible Tables state the number of individuals included in the analysis. |

| | | | | |
|---|---|---|---|---|
| | | (c) *Cohort study*—Summarize follow-up time (e.g., average and total amount) | | Not applicable |
| Outcome data | 15* | *Cohort study*—Report numbers of outcome events or summary measures over time | | Not applicable |
| | | *Case-control study*—Report numbers in each exposure category, or summary measures of exposure | | Not applicable |
| | | *Cross-sectional study*—Report numbers of outcome events or summary measures | Figures and Tables | Data are given as percentages or means. |
| Main results | 16 | (*a*) Give unadjusted estimates and, if applicable, confounder-adjusted estimates and their precision (e.g., 95% confidence interval). Make clear which confounders were adjusted for and why they were included | | Not applicable |
| | | (*b*) Report category boundaries when continuous variables were categorized | | Not applicable |
| | | (*c*) If relevant, consider translating estimates of relative risk into absolute risk for a meaningful time period | | Not applicable |
| Other analyses | 17 | Report other analyses done—e.g. analyses of subgroups and interactions, and sensitivity analyses | Results 5 - 9 | Data from the diagnostic and the *ex vivo* assessments were used for correlation analyses to assess potential interactions. Moreover, re-analyses according to the participants' sex were performed. |
| **Discussion** | | | | |
| Key results | 18 | Summarize key results with reference to study objectives | Discussion 12 | Corresponding information can be found in the first paragraph of the Discussion. |
| Limitations | 19 | Discuss limitations of the study, taking into account sources of potential bias or imprecision. Discuss both direction and magnitude of any potential bias | Discussion 13 - 14 | Potential biases associated with age-related frailty and sex differences in immune subset analysis are discussed:<br>"Although our data are limited in sample size and statistical power for detailed analyses, the consistent association suggests that disrupted bone homeostasis and bone marrow activation alter hematopoiesis in both sexes, […]."<br>"Considering the more rapid loss of trabecular bone after menopause, longitudinal studies are needed to determine how granulocyte and platelet counts, as well as their activation status, change with progressive loss of BMD in both sexes."<br><br>Moreover, limitations of the correlation analyses that might affect the interpretation of the indicated associations are described: "Despite only partially available cardiovascular assessments in our retrospective study, […]."<br><br>Limitations of the *in vitro* approach are discussed in relation to secondary effects of serum mediators that cannot be studied in the current *in vitro* assay:<br>"Other altered serum factors, sCD40L, OPN, SDF-1, showed no effect on SMC calcification, but might contribute to vascular calcification indirectly by exerting effects on endothelial cells, macrophages, or other immune cells that cannot be studied in our current in vitro model." |

| | | | | |
|---|---|---|---|---|
| Interpretation | 20 | Give a cautious overall interpretation of results considering objectives, limitations, multiplicity of analyses, results from similar studies, and other relevant evidence | Discussion 12 - 15 | The identified cellular and molecular signature is cautiously discussed within the context of other studies related to bone health, osteoporosis, vasculature, atherosclerosis, and the bone-vascular cross-talk. Noteworthy findings include:<br><br>1. Alterations in immune cell subsets, ratios and systemic inflammation index:<br>"These findings are consistent with previous reports showing increased neutrophil-to-lymphocyte ratios and systemic inflammation indices in postmenopausal women with low BMD and osteoporotic individuals of both sexes, as well as a negative association between the platelet-to-lymphocyte ratio and lumbar spine BMD in males. In our study, a similar male bias was observed for increased eosinophil and neutrophil counts, whereas platelet counts were increased in both sexes. Although our data are limited in sample size and statistical power for detailed analyses, the consistent association suggests that disrupted bone homeostasis and bone marrow activation alter hematopoiesis in both sexes, with direct vascular consequences. Earlier work has linked low BMD and increased granulocyte counts to clonal hematopoiesis of indeterminate potential (CHIP), a condition that accelerates atherosclerotic processes through clonal expansion of hematopoietic stem cells with somatic mutations. In particular high neutrophil counts have been associated with atherosclerotic cardiovascular disease."<br><br>2. Sex differences in immune cell subpopulations and their potential pathological consequences:<br>"[…] as well as a negative association between the platelet-to-lymphocyte ratio and lumbar spine BMD in males. In our study, a similar male bias was observed for increased eosinophil and neutrophil counts, whereas platelet counts were increased in both sexes."<br>"[…] longitudinal studies are needed to determine how granulocyte and platelet counts, as well as their activation status, change with progressive loss of BMD in both sexes."<br><br>3. *In vitro* and *ex vivo* considerations:<br>While acknowledging the limitations of *in vitro* experiments, our study substantiates the role of PDGF-BB in human smooth muscle cell calcification, as previously established in murine studies. Although we highlight its role as a direct mediator of SMC calcification, *ex vivo* analyses indicates that the pathological bone-vascular cross-talk in osteoporotic individuals is likely driven by dynamic interactions of multiple altered factors in the systemic osteoporotic milieu. We suggest to conceptualize these interactions as a complex adaptive system: "Our analyses rather indicate that the pathological bone-vascular cross-talk needs to be conceptualized as a complex adaptive system (CAS), with simultaneous interactions and potentially dynamic, interdependent cause-and-effect relationships between its components and nested subsystems as outlined in Fig. 6." |
| Generali-zability | 21 | Discuss the generalizability (external validity) of the study results | Discussion 12 - 16 | Findings from this retrospective exploratory study build on previous epidemiological work as well as molecular and cellular insights. In a hypothesis-generating approach, we propose conceptualizing the cross-talk not as driven solely by PDGF-BB, but as a multifactorial interplay within a complex adaptive system, providing a framework for future studies. |

| | | | | |
|---|---|---|---|---|
| **Other information** | | | | |
| Funding | 22 | Give the source of funding and the role of the funders for the present study and, if applicable, for the original study on which the present article is based | Acknow-ledgement 30, 31 | Corresponding information can be found in the Acknowledgement. |

*Give information separately for cases and controls in case-control studies and, if applicable, for exposed and unexposed groups in cohort and cross-sectional studies.

**Note:** An Explanation and Elaboration article discusses each checklist item and gives methodological background and published examples of transparent reporting. The STROBE checklist is best used in conjunction with this article (freely available on the Web sites of PLoS Medicine at http://www.plosmedicine.org/, Annals of Internal Medicine at http://www.annals.org/, and Epidemiology at http://www.epidem.com/). Information on the STROBE Initiative is available at www.strobe-statement.org.
